# Trade-Offs Between Antibacterial Resistance and Fitness Cost in the Production of Metallo-β-Lactamase by Enteric Bacteria Manifest as Sporadic Emergence of Carbapenem Resistance in a Clinical Setting

**DOI:** 10.1101/2020.10.24.353581

**Authors:** Ching Hei Phoebe Cheung, Mohammed Alorabi, Fergus Hamilton, Yuiko Takebayashi, Oliver Mounsey, Kate J. Heesom, Philip B. Williams, O. Martin Williams, Maha Albur, Alasdair P. MacGowan, Matthew B. Avison

## Abstract

Meropenem is a clinically important antibacterial reserved for treatment of multi-resistant infections. In meropenem-resistant bacteria of the family Enterobacteriales, NDM-1 is considerably more common than IMP-1, despite both metallo-β-lactamases (MBLs) hydrolysing meropenem with almost identical kinetics. We show that *bla*_NDM-1_ consistently confers meropenem resistance in wild-type Enterobacteriales, but *bla*_IMP-1_ does not. The reason is higher *bla*_NDM-1_ expression because of its stronger promoter. However, the cost of meropenem resistance is reduced fitness of *bla*_NDM-1_ positive Enterobacteriales because of amino acid starvation. In parallel, from a clinical case, we identified multiple *Enterobacter* spp. isolates carrying a plasmid-encoded *bla*_NDM-1_ having a modified promoter region. This modification lowered MBL production to a level associated with zero fitness cost but, consequently, the isolates were not meropenem resistant. However, we identified a *Klebsiella pneumoniae* isolate from this same clinical case carrying the same *bla*_NDM-1_ plasmid. This isolate was meropenem resistant despite low-level NDM-1 production because of a *ramR* mutation, reducing envelope permeability. Overall, therefore, we show how the resistance/fitness trade-off for MBL carriage can be resolved. The result is sporadic emergence of meropenem resistance in a clinical setting.

## Introduction

β-Lactamases are the most frequent cause of β-lactam resistance among Gram-negative bacteria. In β-lactamases of molecular classes A, C and D, an active site serine catalyses hydrolysis of the β-lactam ring. Members of class B utilize zinc ions in catalysis and are known as metallo-β-lactamases (MBLs). Based on their sequence homology, MBLs are classified into three subclasses: B1, B2 and B3 (1). Chromosomally encoded MBLs belonging to subclasses B2 and B3 have been isolated from environmental and opportunistic pathogenic bacteria such CphA (*Aeromonas hydrophila*) (2), L1 (*Stenotrophomonas maltophilia*) (3), IND (*Chryseobacterium indologenes*) (4), and Sfh-1 (*Serratia fonticola*) (5). However, the most common MBLs in human pathogens are from subclass B1 and are encoded on mobile genetic elements, particularly VIM (6), IMP (7), and NDM (8). These enzymes can efficiently catalyse the hydrolysis of all clinically relevant β-lactams except the monobactams (1).

The genes encoding VIM-1 and IMP-1 are held within class 1 integrons as gene cassettes (6,7). Integrons are gene capture systems consisting of a 5’ conserved sequence including *intI*, encoding an integrase enzyme, an array of gene cassettes, and a 3’ conserved sequence. Gene cassettes are promoter-less and consist of an open reading frame and an adjacent recombination site, *attC*, specifically recognized by the integrase enzyme. A common promoter (Pc) located within the *intI* sequence directs expression of all gene cassettes in an integron (9). There are essentially three strengths of Pc: PcS – strong, PcW – weak, and PcH – intermediate (10).

The *bla*_NDM-1_ gene is not a gene cassette but has been mobilised by an insertion sequence (IS) element, IS*Aba125* (11). This mobilisation also drives expression of *bla*_NDM-1_, because IS*Aba125* carries an outward facing promoter, P_out_ (12).

In a recent UK study, NDM-1 was found to be the dominant MBL in Enterobacteriales clinical isolates, with IMP-1 not being found at all (13). One possible explanation is that NDM-1 is a lipoprotein and has evolved to perform well in the sort of low zinc environment often seen at sites of infection (14), something which is enhanced in various NDM variants, particularly NDM-4 (15). However, it is possible that positive selection for NDM-1 production is driven by something more fundamental. There is some evidence that IMP-1-encoding plasmids only confer borderline resistance to carbapenems in *E. coli* even when zinc concentration are high (e.g. as seen in Ref 16), whereas minimum inhibitory concentrations (MICs) of carbapenems against *E. coli* transconjugants carrying NDM-1 plasmids are much higher (e.g. as seen in Ref 8). We hypothesise that a more consistent ability to confer carbapenem resistance is part of the reason why NDM-1 is dominant over IMP-1. If correct, this would imply that the levels of active enzyme produced are frequently greater for NDM-1-than for IMP-1-positive Enterobacteriales because, catalytically, the enzymes are very similar (8).

The aims of the work presented here was to test the hypothesis that NDM-1 and IMP-1 confer different carbapenem MICs because they are produced at different levels from their native expression environments. Furthermore, we have investigated the fitness trade-offs that come in to play when selection for higher level MBL production is necessary to confer resistance. Finally, we report a clinical case demonstrating how these fitness trade-offs manifest in the real world.

## Results and Discussion

### bla_NDM-1_ is expressed at higher levels than bla_IMP-1_ and confers meropenem resistance in Enterobacteriales clinical isolates

A blastn search of GenBank using the nucleotide sequences of *bla*_IMP-1_ and *bla*_NDM-1_ revealed that, of entries that matched with 100% coverage and identity, *E. coli* (χ^2^=9.82, *p*<0.005) and *Klebsiella* spp. (χ^2^=12.72, *p*<0.0005) are more likely to carry *bla*_NDM-1_ than *bla*_IMP-1_. This analysis is supported by global surveillance data from clinical isolates. For example, from a recent SENTRY study where, of 1298 carbapenem resistant Enterobacteriales analysed in 2014-16, *bla*_NDM_ positivity was 12.7% whilst *bla*_IMP_ positivity was 0.4% (17). In contrast, the non-Enterobacteriales *Pseudomonas* spp. is more likely to carry *bla*_IMP-1_ than *bla*_NDM-1_ (χ^2^=30.18, *p*<0.00001).

We next sought to test the hypothesis that *bla*_NDM-1_ is dominant over *bla*_IMP-1_ in Enterobacteriales because only *bla*_NDM-1_ reliably confers carbapenem resistance. The *bla*_NDM-1_ gene is almost exclusively found downstream of an IS*Aba125* sequence, which provides an outward facing promoter, P_out_, which drives *bla*_NDM-1_ expression (11). In contrast, *bla*_IMP-1_ is encoded as an integron gene cassette (7), and so can be present downstream of several different promoter (Pc) sequences (10). Of the 26 *bla*_IMP-1_ GenBank entries involving *E. coli*, *Klebsiella* spp. and *Enterobacter* spp. where sufficient sequence was present to identify the Pc promoter variant, 24/26 were intermediate strength as previously defined (10) and of these, ten were PcH1 variants (**Table S1**). We therefore chose to compare the impact of carrying *bla*_IMP-1_ located downstream of the PcH1 promoter with *bla*_NDM-1_ located downstream of P_out_ from *ISAba125* on susceptibility to the carbapenem meropenem.

Thirteen out of thirteen *bla*_NDM-1_ Enterobacteriales clinical isolate transformants tested were meropenem resistant, but only 1/13 *bla*_IMP-1_ transformants (**Table S2**). These data support our primary hypothesis, that NDM-1 more readily confers meropenem resistance than IMP-1 in the Enterobacteriales.

IMP-1 and NDM-1 are, in terms of meropenem catalytic efficiency, very similar enzymes (8), so our next hypothesis was that more NDM-1 is produced than IMP-1 in cells, explaining the difference in meropenem MIC. This hypothesis was also supported by experiment; the amount of meropenem hydrolysing activity in cell extracts of representative *bla*_NDM-1_ transformants of *E. coli*, *K. pneumoniae* and *Enterobacter (Klebsiella) aerogenes* was 3 to 6-fold higher than in *bla*_IMP-1_ transformants (*p*<0.002 for each). As expected, elevated meropenem hydrolysing activity was due to greater production of NDM-1 than IMP-1 protein as measured using LC-MS/MS proteomics (**Fig. 1**).

**Figure 1.**
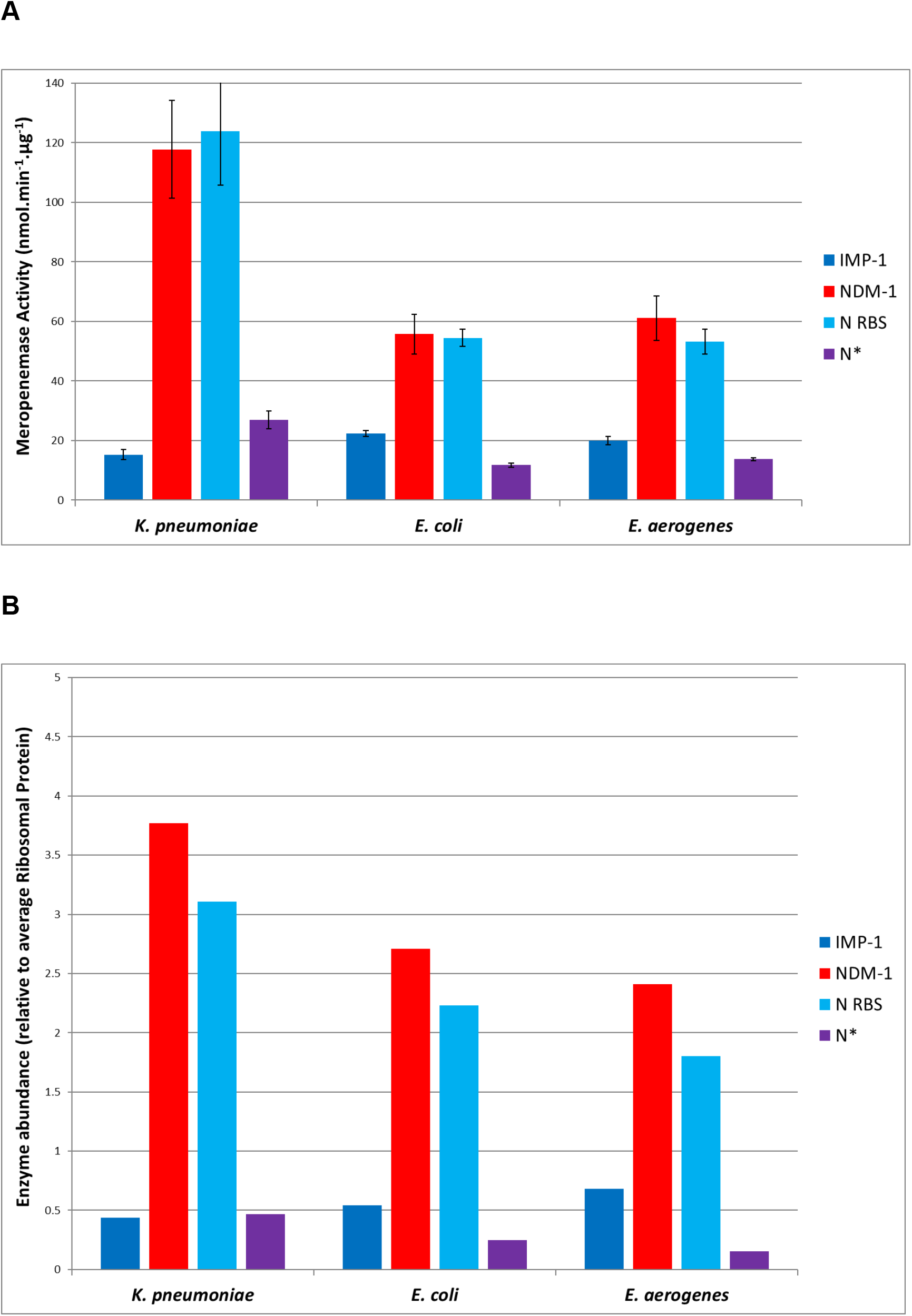
MBL Production in Enterobacteriales carrying *bla*_IMP-1_ or *bla*_NDM-1_ with variant upstream sequences. MBL production was measured in *K. pneumoniae*, *E. coli* or *K. aerogenes* (*Ent. aerogenes*) recombinants carrying the pSU18 cloning vector, into which had been ligated *bla*_IMP-1_ with its upstream Pc(H1) promoter (dark blue bars), *bla*_NDM-1_ with its wild-type IS*Aba125* promoter (bed bars), *bla*_NDM-1_ with site directed mutation to convert its ribosome binding site to be identical to that upstream of *bla*_IMP-1_ (N RBS, light blue bars), and *bla*_NDM-1_ synthesised to have the same upstream sequence as *bla*_IMP-1_ (N*, purple bars). In (**A**) meropenem hydrolysing activity (nmol.min^-1^.mg total protein^-1^) was measured in whole cell extracts. In (**B**) IMP-1 or NDM-1 protein abundance derived from LC-MS/MS analysis of whole cell extracts is reported normalised to the average abundance of 30S and 50S ribosomal proteins in each extract. Data are means +/- Standard Error of the Mean, n=3.

Changing the ribosome binding sequence upstream of *bla*_NDM-1_ to be identical to that found upstream of *bla*_IMP-1_ did not significantly reduce NDM-1 production or meropenem hydrolysing activity. However, generating the N* variant, by replacing the entire *bla*_NDM-1_ upstream sequence with that upstream of *bla*_IMP-1_, reduced NDM-1 production to be very similar to that of IMP-1 in all three species (**Fig. 1**).

### The correlation between high gene expression and fitness cost when carrying bla_NDM-1_ is associated with amino acid starvation

We next investigated whether the greater production of NDM-1 relative to IMP-1 imposes a fitness cost. Using pairwise competition experiments, where transformants were directly competed over 4 days in the absence of β-lactams, we showed that there is no cost of carrying *bla*_IMP-1_ in *E. coli* and *K. pneumoniae*, but there was a significant cost of carrying *bla*_NDM-1_ in both species (**Table 1**).

**Table 1.**
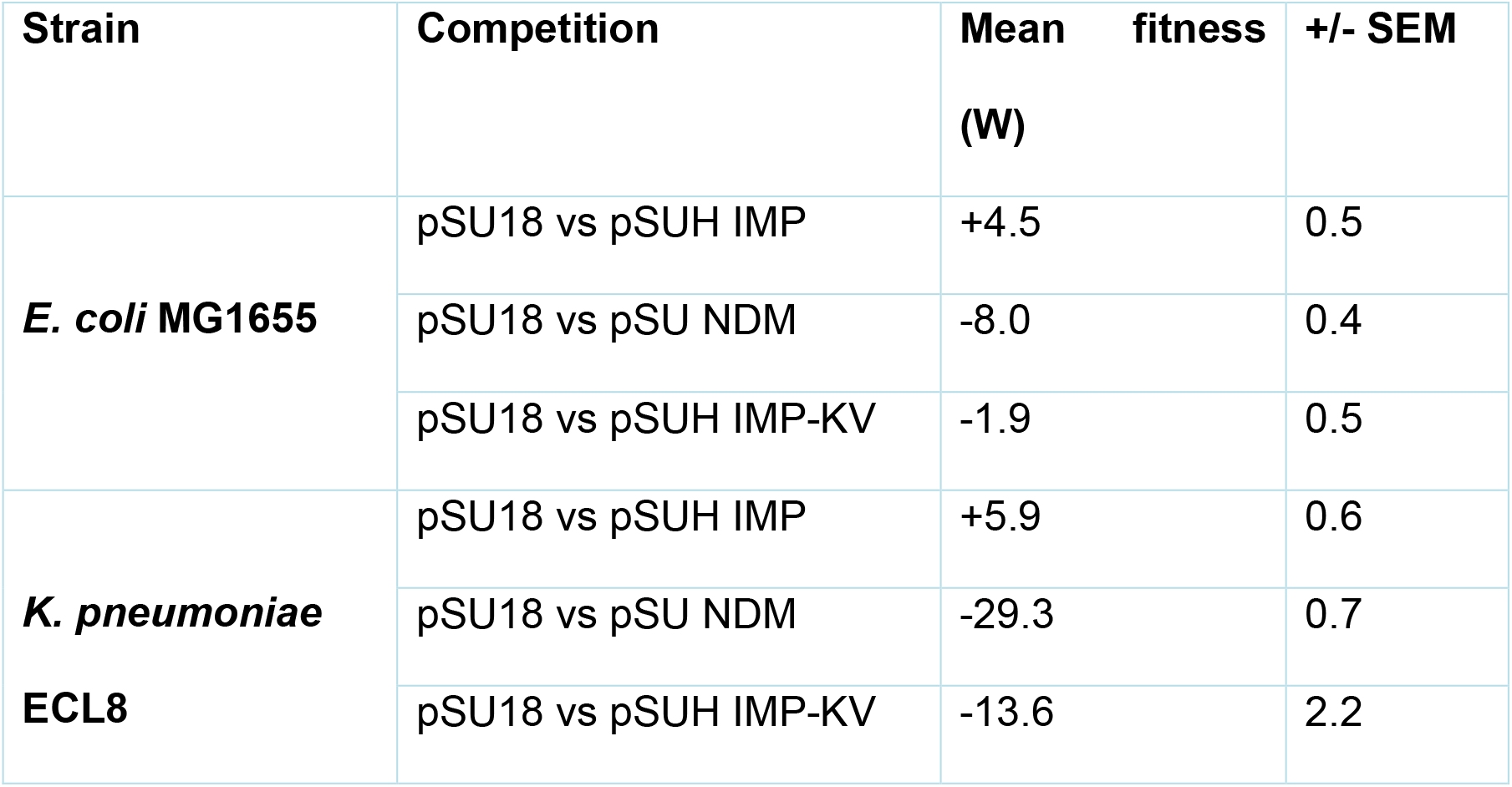
Fitness effect of carrying *bla*_IMP-1_ or *bla*_NDM-1_ in *E. coli* and *K. pneumoniae*.

Higher production of NDM-1 versus IMP-1 could impose a fitness cost because of depletion of resources required to make the additional MBL, or it could be due to some toxicity that the MBL imposes. To differentiate between these possibilities, we investigated the physiological impact of carrying *bla*_IMP-1_ or *bla*_NDM-1_ in *E. coli*. To do this, we used LC-MS/MS proteomics to quantify steady state protein abundance differences in transformants.

Of 1390 proteins identified and quantified in the *bla*_IMP-1_ vs plasmid only control comparison, 66 were significantly up or down regulated (**Table S3**) but Chi squared analysis did not reveal clustering of these proteins into any KEGG functional group, suggesting that there is little concerted physiological response to carrying *bla*_IMP-1_ (**Table S4**). The *bla*_NDM-1_ versus control comparison identified and quantified 1670 proteins, of which 88 were differentially regulated (**Table S5**). In this case Chi squared analysis did identify clustering (**Table S6**) of these regulated proteins into a specific KEGG pathway: eco00260, glycine, serine, and threonine metabolism. Upregulated proteins include the committed enzymes GlyA (18), SerA (19), ThrC (20), and IlvA, which directs these amino acids into other amino acid biosynthetic pathways (21). Therefore, production of NDM-1, which is approximately 6-fold more than production of IMP-1 in *E. coli* (**Fig. 1**), comes with a significantly fitness cost (**Table 1**), which is associated with regulatory signals of amino acid starvation.

### Increasing IMP-1 production increases fitness cost

To further test the hypothesis that the amount of MBL protein production is a major part of the fitness cost imposed by carrying MBL genes and to exclude any NDM-1 specific effects, we aimed to increase IMP-1 production. To do this we turned to our recently reported *bla*_IMP-1_ synonymous lysine codon variant, IMP-1-KV where 17 AAA lysine codons were converted to the alternative synonymous codon, AAG (22). LC-MS/MS proteomics showed that the amount of IMP-1 produced from the variant *bla*_IMP-1_-KV was 2.2-fold (*p*=0.005) more than from wild-type *bla*_IMP-1_ in *E. coli* (**Fig. 2**). As hypothesised, this increase in IMP-1 protein production was associated with an increase in fitness cost, which was approximately 7% per day in *E. coli* and approximately 20% per day in *K. pneumoniae* (p<0.001 for both comparisons) (**Table 1**). We attempted to repeat this experiment by cloning *bla*_IMP-1_ downstream of a strong integron promoter, which drives high-level gene expression, but very few *E. coli* transformants were recovered. In all cases, the transformants had mutations upstream of *bla*_IMP-1_ expected to reduce gene expression, e.g. those affecting the −35 or −10 promoter sequences or the spacing in between. Accordingly, we conclude that the fitness cost of carrying this highly expressed form of *bla*_IMP-1_ is too great for transformants to bear.

**Figure 2.**
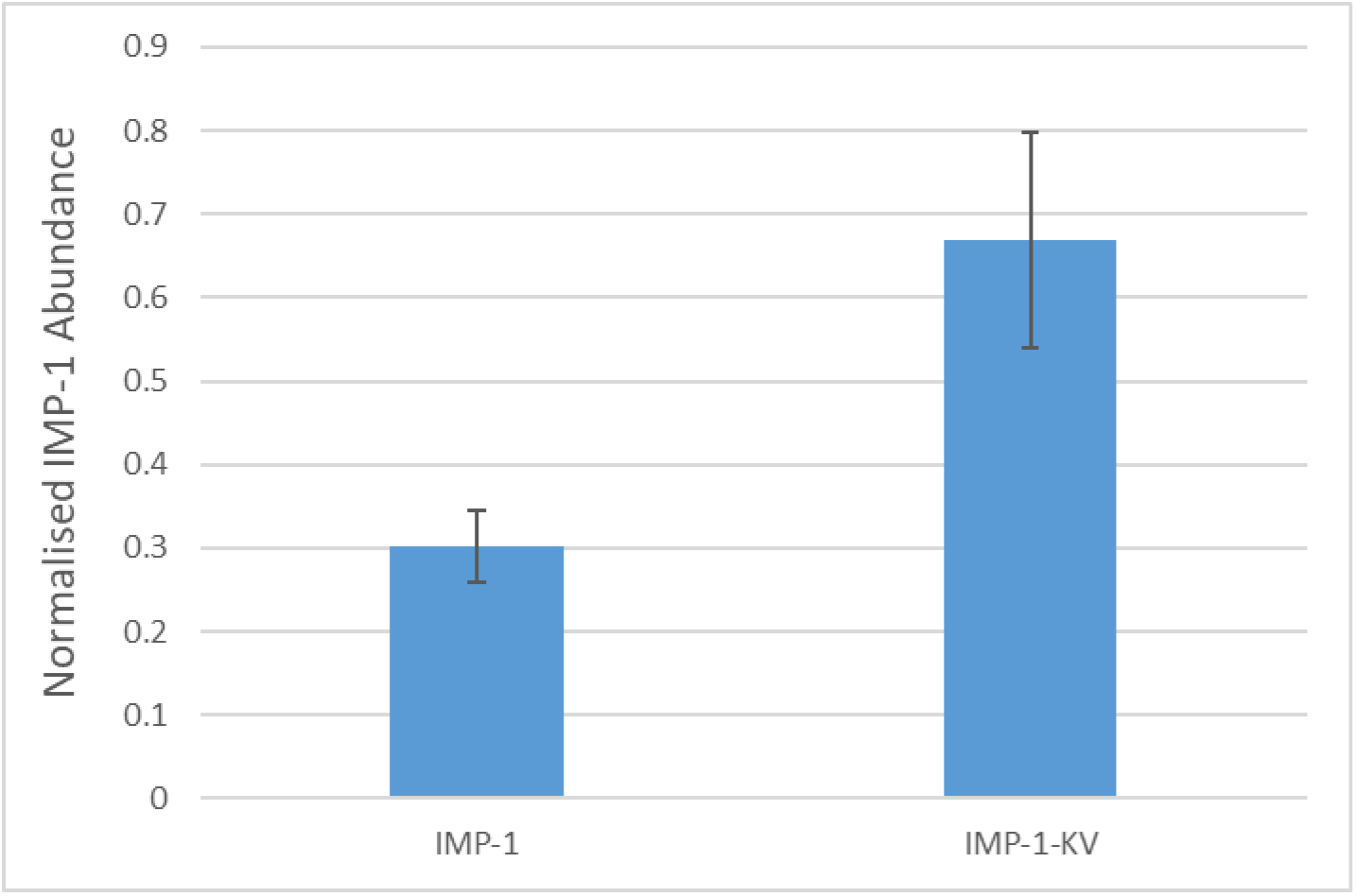
Increased production of IMP-1 following introduction of 17 AAA-AAG lysine codon variants into *bla*_IMP-1_. *E. coli* MG1655 recombinants carry pSU18 with *bla*_IMP-1_ or a variant (22) in which 17 AAA lysine codons had been mutated to AAG (IMP-1-KV) were analysed. IMP-1 protein abundance derived from LC-MS/MS analysis of whole cell extracts is reported normalised to the average abundance of 30S and 50S ribosomal proteins in each extract. Data are means +/- Standard Error of the Mean, n=3.

### Reduced NDM-1 production due to rearrangements in the bla_NDM-1_ promoter region explains lack of meropenem resistance in Enterobacter spp. isolates from a clinical case

A patient was admitted directly to the intensive care unit after developing a small bowel obstruction and an aspiration pneumonia. Bronchoalveolar lavage grew *Citrobacter freundii*, *K. pneumoniae* and *Bacteroides vulgatus*. The patient was initially treated with piperacillin-tazobactam and azithromycin and noted to have a strangulated inguinal hernia which was repaired. Two days after admission, the patient was escalated to meropenem due to continued fever. Vancomycin was added for a possible coagulase negative *Staphylococcus* spp. line infection. They continued to require ventilation and a tracheostomy was performed on day 7. By 20 days after admission, symptoms had resolved and C-reactive protein had fallen to 10 from 368 mg/L on admission, and meropenem was stopped.

Five days later, fever restarted, and a sputum sample grew *K. pneumoniae* resistant to piperacillin-tazobactam and ciprofloxacin, but Extended-Spectrum β-Lactamase (ESBL) negative and susceptible to third-generation cephalosporins. Ceftazidime and vancomycin were started. After 6 days of ceftazidime, a routine multi-resistant coliform screen of the patient’s tracheostomy site noted a ceftazidime resistant *Enterobacter* spp. (Ent1). This was ESBL positive and had a multi-drug resistance phenotype (**Table S7**). Due to an apparently raised meropenem MIC, a Cepheid Xpert-Carba R PCR test was performed, suggesting the presence of *bla*_NDM_. Despite this, Ent1 was not meropenem resistant and so ceftazidime treatment was switched to meropenem. After 10 days of meropenem, the patient improved, and antibiotic therapy was discontinued. Routine screens continued to isolate *Enterobacter* spp. with the same resistance pattern and being *bla*_NDM_ positive (e.g. Ent2) but 12 days after the isolation of Ent1, another routine screen identified an ESBL negative *K. pneumoniae*, which was fully resistant to meropenem (KP3), as well as to third-generation cephalosporins, piperacillin-tazobactam and ciprofloxacin (**Table S7**). The Cepheid Xpert-Carba also identified *bla*_NDM_ in KP3. The patient, however, remained well and continued off antibiotics and was discharged to the surgical ward.

Subsequent routine screens continued to identify this meropenem resistant *K. pneumoniae* and the *bla*_NDM_ positive *Enterobacter* spp. that was not meropenem resistant and specialist infection control precautions were continued.

Whole genome sequence (WGS) analysis of the *Enterobacter* spp. isolates Ent1 and Ent2 showed them to be *Enterobacter hormaechei* and confirmed that *bla*_NDM-1_ is present on the same IncFIB(K) plasmid in both. The plasmid was assembled into a single contig of 84,659 nt carrying genes conferring resistance to amikacin/ciprofloxacin (*aacA4-cr*), rifampicin (*arr*-3), co-trimoxazole (*sul1*) and streptomycin (*aadA1*), all part of the same complex class 1 integron alongside *bla*_NDM-1_. Otherwise, on the chromosome, other relevant resistance genes carried by Ent1 and Ent2 were to ampicillin (*bla*_TEM-1_), and the expected ESBL (*bla*_CTX-M-15_). The isolates also carried chromosomal mutations in *gyrA* (Ser83Ile) and *parC* (Ser80Ile) causing ciprofloxacin resistance. Collectively this acquired resistance genotype explains the antibiograms of Ent1 and Ent2, except for the fact that meropenem resistance should have been provided by the *bla*_NDM-1_ gene but was not.

LC-MS/MS proteomics revealed that NDM-1 production was the same in Ent1 and Ent2. The amount normalised to ribosomal proteins was 0.41 +/- 0.03 (mean +/- SD), which was not significantly different (*p*=0.13) from the amount of IMP-1 produced from its native PcH1 promoter in *bla*_IMP-1_ transformants of *E. coli* and *K. pneumoniae* described above (0.49 +/- 0.18, **Fig. 1**). In contrast, NDM-1 production in Ent1 and Ent2 was significantly different from (*p*<0.0005), and approximately 6-fold less than NDM-1 production in transformants of *E. coli* and *K. pneumoniae* where *bla*_NDM-1_ was expressed from the typical IS*Aba125* P_out_ promoter (3.24 +/- 0.69, **Fig. 1**). This low-level production of NDM-1 in Ent1 and Ent2 likely explains why these isolates are not meropenem resistant (MIC<4 mg/L), as seen for *bla*_IMP-1_ transformants (**Table S2**).

To explain the reason for low-level NDM-1 production in Ent1 and Ent2, we compared the sequence upstream of *bla*_NDM-1_ in these two isolates with those from *E. coli* IR10, the source of the recombinant plasmids used above, and from *K. pneumoniae* KP05-506, which is the original isolate from which *bla*_NDM-1_ was identified (8). We found a significant rearrangement immediately adjacent to the IS*Aba125* P_out_ promoter in Ent1 and Ent2 (**Fig. 3**). There has been an insertion of an element containing a truncated *bla*_OXA-10_ gene.

**Figure 3.**
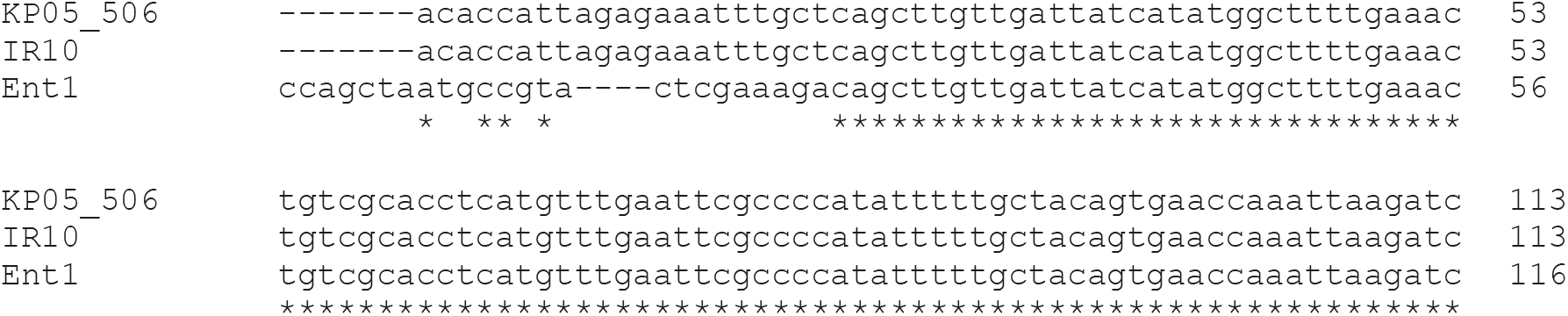
Altered Upstream Sequence in Ent1/2 and KP3 versus *bla*_NDM-1_ Source Sequences. The Clustal Omega alignment used WGS data from two isolates carrying wild-type *bla*_NDM-1_: *E. coli* IR10 and *K. pneumoniae* KP05-506 plus the sequence shared by clinical isolates Ent1, Ent 2 and KP3. Identities across all three sequences are annotated with stars.

The upstream variation seen in Ent1 is rare but not unique. It matched to 14 NCBI database entries reporting isolates collected in China, Taiwan, Japan, Pakistan, and the UK (**Table S8**). Notably, but not commented on by the authors, an *E. coli* transconjugant carrying plasmid pLK78, encoding *bla*_NDM-1_ with this *bla*_OXA-10_ upstream insertion, was not meropenem resistant (23). Moreover, isolates from Pakistan where the *bla*_OXA-10_ insertion upstream of *bla*_NDM-1_ was identified in several related plasmids (24) were originally collected in 2010 and the authors noted that 53% of NDM-1 producing isolates were meropenem susceptible (25).

### Low-level NDM-1 production confers meropenem resistance in a background with reduced envelope permeability

Isolate KP3, from the same clinical case, was meropenem resistant. LC-MS/MS proteomics analysis confirmed that KP3 produced NDM-1 at the same level as Ent1 and Ent2. WGS showed that as well as carrying *bla*_NDM-1_, *aacA4-cr, sul1, arr-3* and *aadA1* on an IncFIB(K) plasmid identical to that found in Ent1 and Ent2, KP3 carried *bla*_TEM-1_ and *bla*_OXA-9_, found together on a second plasmid, plus the chromosomal *bla*_SHV-1_. KP3 also has Ser83Phe and Asp87Ala mutations in GyrA plus a Ser80Ile mutation in ParC explaining ciprofloxacin resistance.

The β-lactamases produced by KP3 in addition to NDM-1 cannot explain the very much higher MIC of meropenem against KP3 versus Ent1 and Ent2. Analysis of KP3 WGS data for known factors that contribute to carbapenem resistance revealed only one: that KP3 is a *ramR* mutant, having an 8 nt insertion into *ramR* after nucleotide 126, causing a frameshift. We have shown that loss of RamR in *K. pneumoniae* leads to enhanced AcrAB-TolC efflux pump production, reduced OmpK35 porin production, and enhanced carbapenem MICs in the presence of weak carbapenemases (26). Hence this mutation in KP3 enhances the meropenem MIC against KP3, making it resistant despite low-level production of NDM-1 due to modification of the IS*Aba125* outward facing promoter region by insertion of a truncated *bla*_OXA-10_.

### Conclusions

Overall, we have observed that modest expression of *bla*_IMP-1_ from a native intermediate strength integron common promoter (PcH1), which is regularly seen in *bla*_IMP-1_ clinical isolates, does not provide meropenem resistance in representative Enterobacteriales strains, but neither does it cause a fitness cost. In contrast, *bla*_NDM-1_ is expressed at higher levels from its native IS*Aba125* outward facing promoter and this gives higher meropenem MICs, clear resistance, but this comes with a significant fitness cost. A fitness cost associated with carrying *bla*_NDM-1_ was also found in a previous report (27). We conclude that the likely reason for this fitness cost, is that NDM-1 is produced at high levels when *bla*_NDM-1_ is expressed from its native promoter, and that this results in amino acid starvation.

Our findings provide a real-world example of fitness/resistance trade-offs. It may be that the reason for *bla*_NDM-1_ being so common in the Enterobacteriales is repeated selective pressure via carbapenem use, driving its presence despite the cost. Alternatively, natural plasmids or certain strains carrying them, or even variant *bla*_NDM_ genes encoded on these plasmids, might have accumulated mutations that compensate for reduced fitness. For example, we have identified the insertion of a truncated *bla*_OXA-10_, damaging the *bla*_NDM-1_ promoter region and reducing NDM-1 production in *Enterobacter* spp. from a clinical case, a genetic arrangement found in commensal carriage Enterobacteriales isolates from as far back as 2010 (25).

Low-level NDM-1 producers avoid the fitness cost associated with *bla*_NDM-1_ carriage but, consequently, are not meropenem resistant. This highlights a potential infection control issue where phenotypic meropenem resistance is necessary for a positive screening outcome. As seen here, the isolates Ent1 and Ent2 were still identified as being of interest due to extra vigilance in respect of a seriously ill patient. With less vigilance, it may have been that the only notice of the presence of an NDM-1 producing isolate in or around this patient would have been following mobilisation of the *bla*_NDM-1_ encoding plasmid into the *ramR* mutant *K. pneumoniae* with reduced envelope permeability, to create meropenem resistant isolate KP3. This ability of reduced envelope permeability to enhance meropenem MIC against a low-level MBL producer may also explain our finding that *bla*_IMP-1_ is more common in *P. aeruginosa*, a species renowned for having much lower envelope permeability than wild-type Enterobacteriales (28). In the context of “under the radar” NDM-1 production defined here, which also relies on reduced envelope permeability, we show that sudden emergence of clinically-relevant meropenem resistance can occur in a manner that is not dependent on new importation events and so cannot be prevented by standard infection control measures.

## Experimental

### Bacteria Used and Susceptibility Testing Assays

Bacterial strains used in the study were *E. coli* MG1655 (29) and a collection of human clinical isolate from urine (a gift from Dr Mandy Wooton, Public Health Laboratory for Wales), a human clinical isolate of *K. aerogenes*, NDM-1 producing isolates of *E. coli* IR10 and *K. pneumoniae* KP05_506 (gifts from Prof T Walsh, Cardiff University), and *K. pneumoniae* strains SM, ECL8 and NCTC 5055 (30). Antibiotic susceptibility was determined using disc testing or broth microdilution MIC assays according to EUCAST guidelines.

### Molecular Biology

Creation of pSUHIMP, being the cloned *bla*_IMP-1_ gene downstream of a native PcH1 was via PCR using template DNA from *P. aeruginosa* clinical isolate 206-3105A (a gift from Dr Mark Toleman, Department of Medical Microbiology, Cardiff University). The sequence of plasmid pYUI-1, the *bla*_IMP-1_ encoding plasmid from this isolate has been deposited under GenBank accession number MH594579. PCR used a forward primer targeting the 5’ end of the PcH1 promoter (5’-ACCCAGTGGACATAAGCCTGTTCGGTTCGTAAACT-3’) and a reverse primer targeting the 5’ end of a *bla*_OXA-1_ gene cassette, which is downstream of *bla*_IMP-1_ in this isolate (5’-AGCGAAGTTGATATGTATTGTG-3’). The PCR amplicon was TA cloned into the pCR2.1TOPO cloning vector (Invitrogen), removed with EcoRI and ligated into EcoRI linearized broad host range p15A-derived vector pSU18 (31). Site directed mutagenesis to create pSUHIMP-KV containing 14 AAA-AAG transitions was performed using the methods and primers previously reported (22). Creation of pSUNDM, being the cloned *bla*_NDM-1_ gene downstream of its native IS*Aba125* promoter in plasmid pSU18 has been reported previously (32). Site directed mutagenesis using pSUNDM as the template was performed using the QuikChange Lightning Site-Directed Mutagenesis Kit (Agilent, UK) according to the manufacturer’s instructions. The aim was to convert the native ribosome binding site upstream of *bla*_NDM-1_ (AAAAGGAAAA<u>CTTG</u>ATGAGCAAGTTATCT) to be the same as that upstream of *bla*_IMP-1_ (AAAAGGAAAA<u>GT</u>ATGAGCAAGTTATCT – differences underlined), using the mutagenic primer 5’-GGGGTTTTTAATGCTGAATAAAAGGAAAAGTATGGAATTGCCCAAT-3’. The resultant plasmid was named pSUNDM-RBS. Switching the entire upstream sequence from the ATG of *bla*_NDM-1_ to be the same as *bla*_IMP-1_ was performed by gene synthesis recreating the entire pSUNDM insert sequence, but with the same upstream sequence carried in pSUHIMP. The resultant plasmid was named pSUNDM-N*

### Proteomic Analysis

A volume of 1 ml of overnight liquid culture was transferred to a 50 ml of fresh LB broth and incubated at 37°C until an OD_600_ of 0.5-0.6 was achieved. Samples were centrifuged at 4,000 rpm for 10 min at 4°C and the supernatants discarded. Cells were re-suspended into lysis buffer (35 ml of 30mM Tris-HCl pH 8) and broken by sonication using a cycle of 1 s on, 1 s off for 3 min at amplitude of 63% using a Sonics Vibracell VC-505TM (Sonics and Materials Inc., Newton, Connecticut, USA). This was followed by centrifugation at 8000 rpm (Sorval RC5B PLUS using an SS-34 rotor) for 15 min at 4°C to pellet non-lysed cells. Soluble proteins were concentrated to a volume of 1 ml using centrifugal filter units (AMICON ULTRA-15, 3 KDa cutoff).

Then, the concentration of the proteins in each sample was measured using Biorad Protein Assay Dye Reagent Concentrate according to the manufacturer’s instructions and normalised. LC-MS/MS was performed and analysed as described previously (33) using 5 μg of protein for each run. Analysis was performed in triplicate, each from a separate batch of cells. Protein abundance was normalised using the average abundance of ribosomal proteins, unless stated in the text.

### Measurement of meropenem hydrolysis

Twenty microlitres of concentrated total cell protein (prepared and assayed for concentration as above) was transferred to 180 μl of 50 mM HEPES (pH 7.5) containing 50 μM ZnSO_4_ and 100 μM meropenem. Change of absorbance was monitored at 299 nm over 10 min. Specific enzyme activity (pmol meropenem hydrolysed per milligram of protein per second) in each extract was calculated using 9600 M^-1^ as the extinction coefficient of meropenem and dividing enzyme activity with the total amount of protein in each assay.

### Pairwise Fitness Cost Experiments

Pairwise competition experiments were performed by using M9 minimal medium to evaluate the fitness cost of carrying pSUHIMP, pSUHIMP-KV or pSUNDM, each relative to the carriage of the pSU18 cloning vector alone. Initially, liquid cultures of both transformants in the pairwise competition were established separately in LB broth at 37°C with shaking at 160 rpm. Then, 5 μl of each overnight liquid culture was inoculated into 10 ml M9 minimal medium separately in flasks and incubated as above for 24 h as before. After this incubation, 5 μl of each overnight M9 minimal medium was again inoculated separately into 10 ml M9 minimal medium as before and grown overnight. The next day, for each competing bacterium, 75 μl of the previous day’s culture was inoculated into fresh 15 ml M9 minimal medium to obtain a mixed culture (day one). After 24 h of incubation, 150 μl of the mixed culture was transferred into a fresh 15 ml M9 minimal medium to obtain the day-two culture. Then, this step was performed successively until the day-four mixed liquid culture was attained. For each pairwise competition experiment, the above process was carried out six times in parallel and on each day, the colony forming units per ml (cfu/ml) of the two bacteria was counted in triplicate using LB agar selective for the cloning vector (the total count of both competitors, as both are chloramphenicol resistant) and agar containing 20 mg/L ceftazidime (to count bacteria producing IMP-1 or NDM-1). The pSU18 containing transformant count was calculated by subtracting the pSUHIMP or pSUNDM containing transformant count from the total count of bacteria in the competition.

The fitness cost of the resistant strain relative to the sensitive strain was estimated by calculating the Malthhusian parameter of the strain (M) as described (34):

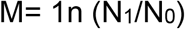

Where N_0_ indicates the density of the strain at the start of the day (cfu/ml) and N_1_ represents the density of the strain at the end of the day (cfu/ml).

Then the selection rate for a pairwise competition is calculated as below:

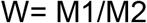

Where M1 represents growth of the sensitive strain and M2 refers to growth of the resistant strain. If R is positive, then M1>M2 which implies that the sensitive strain grows faster than the resistant strain and as a result has a fitness advantage and vice versa.

For each day of competition, 36 values are achieved as for each pair-wise competition there are 6 R values and there are 6 competitions each day (6 mixed cultures a day).

Differences in the two sets of data for each pairwise comparison were assessed using mean and standard deviation of R, and an unpaired t-test (with Welch’s correction) was used to assess the statistical significance of the differences observed.

### Analysis to identify clustering of differentially regulated proteins

The KEGG Mapper tool: http://www.genome.jp/kegg/tool/map_pathway2.html was used. We searched against *E. coli* MG1655 (organism: eco) and entered a list of the Uniprot accession numbers for the differentially regulated proteins. As a control, an equal number of *E. coli* MG1655 Uniprot accession numbers was randomly selected and entered in the KEGG Mapper as above. To determine the total number of proteins in the *E. coli* MG1655 proteome that fall into each KEGG, the entire Uniprot MG1655 accession number list was used to feed the KEGG Mapper tool. These values were used to perform a χ^2^ analysis considering the significance of clustering of differentially regulated proteins by reference to random proteins into a KEGG functional group. To maximise specificity, the comparison with random proteins was performed 10 times, each with a different list of random proteins and the result reported was the lowest χ^2^ value obtained across all 10 comparisons.

### WGS and data analysis

Genomes were sequenced by MicrobesNG (Birmingham, UK) on a HiSeq 2500 instrument (Illumina, San Diego, CA, USA). Reads were trimmed using Trimmomatic (35) and assembled into contigs using SPAdes (36) 3.13.0 (http://cab.spbu.ru/software/spades/) and contigs were annotated using Prokka (37). The presence of plasmids and resistance genes was determined using PlasmidFinder (38) and ResFinder 2.1 (39).

### Ethics Statement

This project is not part of a trial or wider clinical study requiring ethical review. The patient signed to give informed consent that details of their case be referred to in a publication and for educational purposes.

## Acknowledgements

This work was funded by grant MR/N013646/1 to M.B.A., O.M.W., A.P.M. and K.J.H. and grant MR/S004769/1 to M.B.A. from the Antimicrobial Resistance Cross Council Initiative supported by the seven United Kingdom research councils and the National Institute for Health Research, and grant MR/T005408/1 to P.W. and M.B.A. from the Medical Research Council. M. Alorabi. was supported by a Postgraduate Scholarship from the Cultural Bureau of the Kingdom of Saudi Arabia. F.H. was supported by a clinical fellowship from the Wellcome Trust. Genome sequencing was provided by MicrobesNG (http://www.microbesng.uk). We are grateful to Dr Aisha Alamri and to Ka Wang Mak, both lately of the School of Cellular & Molecular Medicine, University of Bristol for constructing pSUHIMP and attempting to clone *bla*_IMP-1_ downstream of PcS.

**The authors declare no conflicts of interest.**

## Author Contributions

Conceived the Study: M.B.A., F.H.

Collection of Data: C.C., M. Alorabi, Y.T., O.M, K.J.H, F.H., supervised by M. Albur, A.P.M., M.B.A.

Cleaning and Analysis of Data: C.C., M. Alorabi, Y.T., O.M, K.J.H, F.H., O.M.W., P. W. supervised by M. Albur, A.P.M., M.B.A.

Initial Drafting of Manuscript: M. Alorabi, F.H., M.B.A.

Corrected and Approved Manuscript: All Authors.

