## Supplementary Data for "Trade-Offs Between Antibacterial Resistance and Fitness Cost in the Production of Metallo-β-Lactamase by Enteric Bacteria Manifest as Sporadic Emergence of Carbapenem Resistance in a Clinical Setting"

**Table S1. Integron promoter types associated with *bla*$_{IMP-1}$ containing integrons in *Klebsiella* spp., *Enterobacter* spp. and *E. coli*.**

| Host Species | Accession (Reference) | Promoter Sequence | Type | Strength |
|---|---|---|---|---|
| *E. coli* | AB733642.1 Unpublished | TGGACATAAGCCTGTTCGGTTCGTAAACT | PcH1 | Intermediate |
| *Klebsiella* spp. | AP018455.1 Unpublished | TTGACATAAGCCTGTTCGGTTCGTAAGCT | PcH2 | Intermediate |
| *Klebsiella* spp. | AP018454.1 Unpublished | TTGACATAAGCCTGTTCGGTTCGTAAGCT | PcH2 | Intermediate |
| *Klebsiella* spp. | LC169566.1 Ref S1 | TGGACATAAGCCTGTTCGGTTCGTAAACT | PcH1 | Intermediate |
| *Klebsiella* spp. | KC200566.1 Ref S2 | TTGACATAAGCCTGTTCGGTTCGTAAACT | PcS | Strong |
| *Klebsiella* spp. | AB469046.1 Ref S3 | TGGACATAAGCCTGTTCGGTTGGTAAGCT | PcW-TGN10 | Intermediate |
| *Klebsiella* spp. | D29636.1 Unpublished | TGGACATAAGCCTGTTCGGTTCGTAAACT | PcH1 | Intermediate |
| *Klebsiella* spp. | AP022349.1 Ref S4 | TGGACATAAGCCTGTTCGGTTCGTAAACT | PcH1 | Intermediate |
| *Klebsiella* spp. | AP022081.1 Unpublished | TGGACATAAGCCTGTTCGGTTCGTAAACT | PcH1 | Intermediate |
| *Klebsiella spp.* | AP018557.1 Ref S5 | TGGACATAAGCCTGTTCGGTTCGTAAACT | PcH1 | Intermediate |
| *Enterobacter* spp. | AP018352.1 Ref S6 | TGGACATAAGCCTGTTCGGTTGGTAAGCT | PcW-TGN10 | Intermediate |
| *Enterobacter* spp. | AP018351.1 Ref S6 | TGGACATAAGCCTGTTCGGTTCGTAAACT | PcH1 | Intermediate |
| *Enterobacter* spp. | LC310853.1 | TGGACATAAGCCTGTTCGGTTCGTAAACT | PcH1 | Intermediate |

| | | | | |
|---|---|---|---|---|
| | Ref S6 | | | |
| *Enterobacter* spp. | AP019382.1<br>Ref S7 | <u>TGGACA</u>TAAGCCTGTTCGGTTCG<u>TAAACT</u> | PcH1 | Intermediate |
| *Enterobacter* spp. | AP018350.1<br>Ref S6 | <u>TGGACA</u>TAAGCCTGTTCGGTTCG<u>TAAACT</u> | PcH1 | Intermediate |
| *Enterobacter* spp. | AP019384.1<br>Ref S7 | <u>TTGACA</u>TAAGCCTGTTCGGTTCG<u>TAAACT</u> | PcS | Strong |
| *Enterobacter* spp. | KC675185.1<br>Unpublished | <u>TGGACA</u>TAAGCCTGTTCGGTT<u>G</u>G<u>TAAGCT</u> | PcW-TGN10 | Intermediate |
| *Enterobacter* spp. | AP022129.1<br>Unpublished | <u>TGGACA</u>TAAGCCTGTTCGGTT<u>G</u>G<u>TAAGCT</u> | PcW-TGN10 | Intermediate |
| *Enterobacter* spp. | LC508022.1<br>Unpublished | <u>TGGACA</u>TAAGCCTGTTCGGTT<u>G</u>G<u>TAAGCT</u> | PcW-TGN10 | Intermediate |
| *Enterobacter* spp. | AP019383.1<br>Ref S7 | <u>TGGACA</u>TAAGCCTGTTCGGTT<u>G</u>G<u>TAAGCT</u> | PcW-TGN10 | Intermediate |
| *Enterobacter* spp. | AP019386.1<br>Ref S7 | <u>TGGACA</u>TAAGCCTGTTCGGTT<u>G</u>G<u>TAAGCT</u> | PcW-TGN10 | Intermediate |
| *Enterobacter* spp. | AP019387.1<br>Ref S7 | <u>TGGACA</u>TAAGCCTGTTCGGTT<u>G</u>G<u>TAAGCT</u> | PcW-TGN10 | Intermediate |
| *Enterobacter* spp. | AP019388.1<br>Ref S7 | <u>TGGACA</u>TAAGCCTGTTCGGTT<u>G</u>G<u>TAAGCT</u> | PcW-TGN10 | Intermediate |
| *Enterobacter* spp. | AP022628.1<br>Ref S8 | <u>TGGACA</u>TAAGCCTGTTCGGTT<u>G</u>G<u>TAAGCT</u> | PcW-TGN10 | Intermediate |
| *Enterobacter* spp. | LC532225.1<br>Ref S8 | <u>TGGACA</u>TAAGCCTGTTCGGTT<u>G</u>G<u>TAAGCT</u> | PcW-TGN10 | Intermediate |
| *Enterobacter* spp. | LC532227.1<br>Ref S8 | <u>TGGACA</u>TAAGCCTGTTCGGTT<u>G</u>G<u>TAAGCT</u> | PcW-TGN10 | Intermediate |

**Table S2. Meropenem susceptibility testing for transformants carrying *bla*$_{\text{IMP-1}}$ or *bla*$_{\text{NDM-1}}$ encoding recombinant plasmids**

| Bacterium | IMP-1 | NDM-1 |
|---|---|---|
| *E. coli* MG1655 | 18 | 11 |
| *E. coli* 3 | 18 | 6 |
| *E. coli* 4 | 18 | 6 |
| *E. coli* 6 | 13 | 6 |
| *E. coli* 10 | 17 | 6 |
| *E. coli* 13 | 20 | 12 |
| *E. coli* 14 | 18 | 11 |
| *E. coli* 16 | 22 | 10 |
| *E. coli* 17 | 18 | 10 |
| *K. aerogenes* | 18 | 10 |
| *K. pnuemoniae* SM | 20 | 10 |
| *K. pneumoniae* ECL8 | 22 | 14 |
| *K. pneumoniae* NCTC 5055 | 18 | 6 |

Data reported are inhibition zone diameters (mm) around a meropenem disc. Data are averages of three repetitions rounded to the nearest integer. Highlights in Red are resistant (<16 mm) based on EUCAST breakpoints. Green are susceptible (≥22 mm). Orange are intermediate (21-16 mm). The MICs of meropenem against *E. coli* 16 and *K. pneumoniae* ECL8 were 1 mg/L when carrying *bla*$_{\text{IMP-1}}$ and >64 mg/L when carrying *bla*$_{\text{NDM-1}}$.

**Table S3. Proteins with significant abundance changes in *E. coli* carrying pSU18::*bla*$_{IMP-1}$ versus plasmid only control**

Data provided are raw protein abundance values for three biological replicates (E1-E3). Green, upregulated; red, downregulated > 2 fold.

| Accession | Description | pSU E1 | pSU E2 | pSU E3 | IMP E1 | IMP E2 | IMP E3 | T-Test | Fold Change |
|---|---|---|---|---|---|---|---|---|---|
| P52699 | Beta-lactamase IMP-1 | | | | 1.16E+10 | 3.49E+09 | 2.36E+09 | <0.001 | >20.00 |
| P0A6A0 | Probable protein kinase UbiB | | | | 1.36E+07 | 4.75E+06 | 4.59E+06 | <0.001 | >20.00 |
| P0A725 | UDP-3-O-[3-hydroxymyristoyl] N-acetylglucosamine deacetylase | | | | 8.30E+07 | 1.19E+07 | 1.25E+08 | <0.001 | >20.00 |
| P0A7Y4 | Ribonuclease H | | | | 1.82E+07 | 5.29E+06 | 6.20E+06 | <0.001 | >20.00 |
| P0A9I3 | Glycine cleavage system transcriptional repressor | | | | 6.57E+07 | 1.41E+08 | 1.13E+08 | <0.001 | >20.00 |
| P0AAB8 | Universal stress protein D | | | | 8.17E+07 | 4.30E+07 | 3.21E+07 | <0.001 | >20.00 |
| P0AAS0 | Inner membrane protein YlaC | | | | 5.76E+07 | 1.97E+07 | 2.07E+07 | <0.001 | >20.00 |
| P0AB26 | Uncharacterized lipoprotein YceB | | | | 3.56E+07 | 5.96E+06 | 6.23E+06 | <0.001 | >20.00 |
| P0ABE2 | Protein BolA | | | | 6.68E+08 | 2.71E+07 | 2.05E+07 | <0.001 | >20.00 |
| P0ACU2 | HTH-type transcriptional regulator RutR | | | | 1.56E+07 | 9.10E+06 | 3.09E+06 | <0.001 | >20.00 |
| P0AF56 | Uncharacterized protein YjcO | | | | 2.02E+07 | 6.95E+06 | 1.54E+07 | <0.001 | >20.00 |
| P0AFX0 | Ribosome hibernation promoting factor | | | | 2.01E+07 | 3.32E+07 | 5.35E+07 | <0.001 | >20.00 |
| P0AFX4 | Regulator of sigma D | | | | 3.69E+07 | 1.69E+07 | 1.60E+07 | <0.001 | >20.00 |
| P0AG03 | 3-octaprenyl-4-hydroxybenzoate carboxy-lyase partner protein | | | | 1.56E+07 | 5.01E+06 | 4.33E+06 | <0.001 | >20.00 |
| P0AGL5 | Ribosome association toxin RatA | | | | 1.24E+08 | 8.17E+06 | 1.16E+07 | <0.001 | >20.00 |
| P13445 | RNA polymerase sigma factor RpoS | | | | 6.36E+07 | 4.67E+07 | 6.57E+06 | <0.001 | >20.00 |
| P17445 | NAD/NADP-dependent betaine aldehyde dehydrogenase | | | | 6.74E+07 | 3.81E+07 | 2.58E+07 | <0.001 | >20.00 |
| P21367 | Uncharacterized protein YcaC | | | | 2.16E+07 | 5.59E+07 | 1.22E+07 | <0.001 | >20.00 |
| P36649 | Blue copper oxidase CueO | | | | 3.98E+07 | 2.08E+07 | 1.74E+07 | <0.001 | >20.00 |
| P36659 | Curved DNA-binding protein | | | | 2.99E+07 | 1.03E+07 | 9.95E+06 | <0.001 | >20.00 |
| P37757 | Uncharacterized isomerase YddE | | | | 4.34E+07 | 7.70E+06 | 6.18E+06 | <0.001 | >20.00 |
| P39334 | HTH-type transcriptional repressor BdcR | | | | 1.20E+07 | 3.95E+07 | 4.94E+06 | <0.001 | >20.00 |

| | | | | | | | | | |
|---|---|---|---|---|---|---|---|---|---|
| P75691 | Uncharacterized zinc-type alcohol dehydrogenase-like protein YahK | | | | 3.68E+07 | 4.25E+07 | 4.44E+07 | <0.001 | >20.00 |
| P75767 | Putative gluconeogenesis factor | | | | 2.34E+07 | 8.20E+06 | 4.59E+06 | <0.001 | >20.00 |
| P31658 | Molecular chaperone Hsp31 and glyoxalase 3 | | 1.73E+07 | 3.09E+07 | 1.32E+08 | 1.57E+08 | 1.56E+08 | 0.001 | 6.14 |
| P08997 | Malate synthase A | 1.53E+08 | 1.53E+08 | 3.93E+07 | 4.84E+08 | 6.81E+08 | 8.96E+08 | 0.005 | 5.97 |
| P0A9G6 | Isocitrate lyase | 1.98E+08 | 7.19E+08 | 2.20E+08 | 1.25E+09 | 2.52E+09 | 2.42E+09 | 0.010 | 5.44 |
| P23843 | Periplasmic oligopeptide-binding protein | 1.97E+08 | 7.52E+07 | 1.03E+07 | 6.65E+08 | 4.20E+08 | 2.97E+08 | 0.019 | 4.90 |
| P0AF98 | Lipopolysaccharide export system permease protein LptF | 1.12E+07 | | 5.87E+06 | 4.34E+07 | | 3.80E+07 | 0.007 | 4.78 |
| P0CK95 | Putative lipoprotein AcfD homolog | 2.98E+08 | | 9.46E+07 | 1.28E+09 | 7.81E+08 | 5.95E+08 | 0.044 | 4.51 |
| P37666 | Glyoxylate/hydroxypyruvate reductase B | 8.95E+07 | 5.88E+07 | 3.50E+07 | 3.51E+08 | 2.33E+08 | 1.72E+08 | 0.013 | 4.12 |
| P75913 | Glyoxylate/hydroxypyruvate reductase A | 2.22E+07 | 1.91E+07 | 1.68E+07 | 1.14E+08 | 7.94E+07 | 3.80E+07 | 0.029 | 3.99 |
| P04693 | Aromatic-amino-acid aminotransferase | 7.50E+07 | 5.17E+07 | 3.13E+07 | 2.27E+08 | 6.93E+07 | 2.80E+08 | 0.048 | 3.65 |
| P08200 | Isocitrate dehydrogenase [NADP] | 1.17E+09 | 9.98E+08 | 9.08E+08 | 4.01E+09 | 3.30E+09 | 3.05E+09 | 0.001 | 3.37 |
| P60560 | GMP reductase | 5.16E+07 | 2.68E+07 | 2.25E+07 | 1.64E+08 | 1.07E+08 | 4.99E+07 | 0.049 | 3.18 |
| P69910 | Glutamate decarboxylase beta | 1.96E+08 | 1.92E+07 | 1.00E+07 | 1.57E+08 | 2.52E+08 | 2.88E+08 | 0.047 | 3.10 |
| P27306 | Soluble pyridine nucleotide transhydrogenase | 6.82E+07 | 6.64E+07 | 2.84E+07 | 1.75E+08 | 1.60E+08 | 1.67E+08 | 0.001 | 3.08 |
| P26648 | Cell division protein FtsP | 3.62E+07 | 5.28E+06 | 6.53E+06 | 6.42E+07 | 5.07E+07 | 3.28E+07 | 0.035 | 3.08 |
| P07650 | Thymidine phosphorylase | 2.17E+07 | 4.39E+07 | 3.31E+07 | 7.70E+07 | 1.00E+08 | 1.21E+08 | 0.005 | 3.02 |
| P0AET8 | 7-alpha-hydroxysteroid dehydrogenase | 5.60E+07 | 4.23E+07 | 3.44E+07 | 1.49E+08 | 1.41E+08 | 9.74E+07 | 0.004 | 2.92 |
| P0AFG6 | Dihydrolipoyllysine-residue succinyltransferase component of 2-oxoglutarate dehydrogenase complex | 2.92E+08 | 3.20E+08 | 1.02E+08 | 5.87E+08 | 6.30E+08 | 8.64E+08 | 0.007 | 2.92 |
| P0AC59 | Glutaredoxin-2 | 6.43E+07 | 2.29E+07 | 1.97E+07 | 1.47E+08 | 1.03E+08 | 6.00E+07 | 0.040 | 2.89 |
| P12758 | Uridine phosphorylase | 4.66E+08 | 3.24E+08 | 2.99E+08 | 9.77E+08 | 1.20E+09 | 8.26E+08 | 0.003 | 2.76 |
| P61889 | Malate dehydrogenase | 7.47E+08 | 1.39E+09 | 1.33E+09 | 3.99E+09 | 3.17E+09 | 2.38E+09 | 0.008 | 2.75 |
| P39160 | D-mannonate oxidoreductase | 2.84E+07 | 4.46E+07 | 1.05E+07 | 5.75E+07 | 9.79E+07 | 6.85E+07 | 0.020 | 2.68 |
| P00934 | Threonine synthase | 9.00E+07 | 5.73E+07 | 4.66E+07 | 2.28E+08 | 1.20E+08 | 1.57E+08 | 0.020 | 2.60 |
| P0A6J5 | D-amino acid dehydrogenase small subunit | | 7.86E+07 | 5.81E+07 | 2.20E+08 | 1.33E+08 | 1.75E+08 | 0.024 | 2.57 |
| P0A6K6 | Phosphopentomutase | 1.81E+08 | 1.03E+08 | 7.32E+07 | 2.55E+08 | 2.85E+08 | 3.31E+08 | 0.006 | 2.44 |
| P0C0L2 | PeroxiredoxinmC | 2.11E+07 | 2.40E+07 | | 4.26E+07 | 6.83E+07 | 5.41E+07 | 0.022 | 2.44 |

| | | | | | | | | | |
|---|---|---|---|---|---|---|---|---|---|
| P0ABH7 | Citrate synthase | 1.43E+08 | 9.03E+08 | 6.39E+08 | 1.44E+09 | 1.14E+09 | 1.52E+09 | 0.016 | 2.44 |
| P0A991 | Fructose-bisphosphate aldolase class 1 | 4.31E+07 | 6.64E+07 | 6.56E+06 | 1.02E+08 | 8.37E+07 | 9.03E+07 | 0.021 | 2.38 |
| P32695 | tRNA-dihydrouridine synthase A | | 7.20E+06 | 9.85E+06 | | 1.81E+07 | 2.16E+07 | 0.017 | 2.33 |
| P00370 | NADP-specific glutamate dehydrogenase | 1.05E+07 | 2.18E+07 | 1.71E+07 | 3.05E+07 | 4.57E+07 | 3.67E+07 | 0.009 | 2.28 |
| Q46845 | Disulfide-bond oxidoreductase YghU | 1.61E+07 | 1.66E+07 | 1.95E+07 | 5.27E+07 | 3.84E+07 | 2.80E+07 | 0.018 | 2.28 |
| P77318 | Uncharacterized sulfatase YdeN | | 1.14E+07 | 6.55E+06 | | 1.94E+07 | 2.15E+07 | 0.024 | 2.28 |
| P0AC33 | Fumarate hydratase class I, aerobic | 1.12E+08 | 2.29E+08 | 1.62E+08 | 4.14E+08 | 2.15E+08 | 4.58E+08 | 0.038 | 2.17 |
| P00509 | Aspartate aminotransferase | 3.29E+08 | 2.31E+08 | 2.53E+08 | 8.28E+08 | 4.02E+08 | 5.25E+08 | 0.037 | 2.16 |
| P15034 | Xaa-Pro aminopeptidase | 1.12E+08 | 4.34E+07 | 2.65E+07 | 1.29E+08 | 1.19E+08 | 1.42E+08 | 0.032 | 2.14 |
| P14407 | Fumarate hydratase class I, anaerobic | 1.21E+08 | 1.21E+08 | 8.64E+07 | 2.53E+08 | 1.74E+08 | 2.64E+08 | 0.009 | 2.10 |
| P0A968 | Cold shock-like protein CspD | | 7.04E+07 | 8.87E+07 | 3.26E+07 | | 4.17E+07 | 0.027 | 0.47 |
| P0A6S0 | Flagellar L-ring protein | 5.77E+07 | 4.19E+07 | 5.62E+07 | 2.92E+07 | 1.35E+07 | 2.59E+07 | 0.007 | 0.44 |
| P76010 | Flagellar brake protein YcgR | 8.38E+07 | 3.77E+07 | 7.19E+07 | 4.53E+07 | 2.30E+07 | 1.65E+07 | 0.045 | 0.44 |
| P05707 | Sorbitol-6-phosphate 2-dehydrogenase | | 4.14E+08 | 4.56E+08 | | 1.25E+08 | 1.85E+08 | 0.008 | 0.36 |
| P0AC30 | Cell division protein FtsX | 5.30E+07 | | 4.39E+07 | 2.94E+07 | 6.01E+06 | 9.75E+06 | 0.022 | 0.31 |
| P77433 | Uncharacterized protein YkgG | 1.94E+07 | 1.57E+07 | 3.46E+07 | 4.39E+06 | 3.15E+06 | 1.06E+07 | 0.025 | 0.26 |
| P76272 | Uncharacterized protein YebT | 7.53E+06 | 1.29E+07 | 1.59E+06 | | | | <0.001 | <0.05 |
| P76440 | NAD-dependent dihydropyrimidine dehydrogenase subunit PreT | 1.14E+07 | 9.76E+06 | 4.43E+06 | | | | <0.001 | <0.05 |

**Table S4. Chi Squared analysis of KEGG functional grouping of proteins with significant abundance changes in *E. coli* carrying pSU18::*bla*$_{IMP-1}$ versus plasmid only control.**

| KEGG | Total in Genome | Number with abundance change | random | Chi Sq |
|---|---|---|---|---|
| eco00010 Glycolysis / Gluconeogenesis - Escherichia coli K-12 MG1655 | 43 | 1 | 0 | 1.01 |
| eco00020 Citrate cycle | 27 | 6 | 2 | 2.35 |
| eco00030 Pentose phosphate pathway - Escherichia coli K-12 MG1655 | 30 | 1 | 0 | 1.02 |
| eco00040 Pentose and glucuronate interconversions - Escherichia coli K-12 MG1655 | 27 | 1 | 0 | 1.02 |
| eco00051 Fructose and mannose metabolism - Escherichia coli K-12 MG1655 | 41 | 1 | 0 | 1.01 |
| eco00052 Galactose metabolism - Escherichia coli K-12 MG1655 | 36 | 0 | 0 | - |
| eco00130 Ubiquinone and other terpenoid-quinone biosynthesis - Escherichia coli K-12 MG1655 | 20 | 1 | 1 | 0 |
| eco00190 Oxidative phosphorylation - Escherichia coli K-12 MG1655 | 41 | 0 | 0 | - |
| eco00220 Arginine biosynthesis - Escherichia coli K-12 MG1655 | 18 | 2 | 0 | 2.12 |
| eco00230 Purine metabolism - Escherichia coli K-12 MG1655 | 89 | 2 | 2 | 0 |
| eco00240 Pyrimidine metabolism - Escherichia coli K-12 MG1655 | 65 | 2 | 3 | 0.21 |
| eco00250 Alanine, aspartate and glutamate metabolism - Escherichia coli K-12 MG1655 | 31 | 3 | 0 | 3.15 |
| eco00260 Glycine, serine and threonine metabolism - Escherichia coli K-12 MG1655 | 36 | 3 | 1 | 1.06 |
| eco00261 Monobactam biosynthesis - Escherichia coli K-12 MG1655 | 10 | 0 | 1 | 1.05 |
| eco00270 Cysteine and methionine metabolism - Escherichia coli K-12 MG1655 | 32 | 3 | 1 | 1.07 |
| eco00280 Valine, leucine and isoleucine degradation - Escherichia coli K-12 MG1655 | 11 | 0 | 0 | - |
| eco00290 Valine, leucine and isoleucine biosynthesis - Escherichia coli K-12 MG1655 | 16 | 0 | 0 | - |
| eco00300 Lysine biosynthesis - Escherichia coli K-12 MG1655 | 15 | 0 | 1 | 1.03 |
| eco00310 Lysine degradation - Escherichia coli K-12 MG1655 | 12 | 1 | 0 | 1.04 |
| eco00330 Arginine and proline metabolism - Escherichia coli K-12 MG1655 | 25 | 1 | 0 | 1.02 |
| eco00350 Tyrosine metabolism - Escherichia coli K-12 MG1655 | 9 | 2 | 0 | 2.25 |

| | | | | |
|---|---|---|---|---|
| eco00360 Phenylalanine metabolism - Escherichia coli K-12 MG1655 | 30 | 3 | 0 | 3.16 |
| eco00380 Tryptophan metabolism - Escherichia coli K-12 MG1655 | 9 | 0 | 0 | - |
| eco00400 Phenylalanine, tyrosine and tryptophan biosynthesis - Escherichia coli K-12 MG1655 | 21 | 2 | 0 | 2.10 |
| eco00401 Novobiocin biosynthesis - Escherichia coli K-12 MG1655 | 4 | 2 | 0 | 2.67 |
| eco00410 beta-Alanine metabolism - Escherichia coli K-12 MG1655 | 15 | 1 | 0 | 1.03 |
| eco00430 Taurine and hypotaurine metabolism - Escherichia coli K-12 MG1655 | 6 | 1 | 0 | 1.09 |
| eco00460 Cyanoamino acid metabolism - Escherichia coli K-12 MG1655 | 6 | 0 | 0 | - |
| eco00473 D-Alanine metabolism - Escherichia coli K-12 MG1655 | 4 | 0 | 0 | - |
| eco00480 Glutathione metabolism - Escherichia coli K-12 MG1655 | 19 | 1 | 0 | 1.03 |
| eco00520 Amino sugar and nucleotide sugar metabolism - Escherichia coli K-12 MG1655 | 46 | 0 | 1 | 1.01 |
| eco00521 Streptomycin biosynthesis - Escherichia coli K-12 MG1655 | 9 | 0 | 0 | - |
| eco00523 Polyketide sugar unit biosynthesis - Escherichia coli K-12 MG1655 | 6 | 0 | 0 | - |
| eco00525 Acarbose and validamycin biosynthesis - Escherichia coli K-12 MG1655 | 4 | 0 | 0 | - |
| eco00540 Lipopolysaccharide biosynthesis - Escherichia coli K-12 MG1655 | 31 | 1 | 0 | 1.02 |
| eco00550 Peptidoglycan biosynthesis - Escherichia coli K-12 MG1655 | 23 | 0 | 1 | 1.02 |
| eco00561 Glycerolipid metabolism - Escherichia coli K-12 MG1655 | 12 | 0 | 0 | - |
| eco00564 Glycerophospholipid metabolism - Escherichia coli K-12 MG1655 | 30 | 0 | 0 | - |
| eco00620 Pyruvate metabolism - Escherichia coli K-12 MG1655 | 51 | 6 | 1 | 3.84 |
| eco00627 Aminobenzoate degradation - Escherichia coli K-12 MG1655 | 8 | 0 | 0 | - |
| eco00630 Glyoxylate and dicarboxylate metabolism - Escherichia coli K-12 MG1655 | 41 | 5 | 3 | 0.55 |
| eco00640 Propanoate metabolism - Escherichia coli K-12 MG1655 | 38 | 0 | 0 | - |
| eco00650 Butanoate metabolism - Escherichia coli K-12 MG1655 | 37 | 1 | 0 | 1.01 |
| eco00660 C5-Branched dibasic acid metabolism - Escherichia coli K-12 MG1655 | 10 | 0 | 0 | - |
| eco00680 Methane metabolism - Escherichia coli K-12 MG1655 | 27 | 2 | 0 | 2.08 |
| eco00740 Riboflavin metabolism - Escherichia coli K-12 MG1655 | 8 | 0 | 0 | - |
| eco00750 Vitamin B6 metabolism - Escherichia coli K-12 MG1655 | 9 | 1 | 0 | 1.06 |

| | | | | |
|---|---|---|---|---|
| eco00760 Nicotinate and nicotinamide metabolism - Escherichia coli K-12 MG1655 | 22 | 1 | 3 | 1.10 |
| eco00770 Pantothenate and CoA biosynthesis - Escherichia coli K-12 MG1655 | 23 | 0 | 0 | - |
| eco00790 Folate biosynthesis - Escherichia coli K-12 MG1655 | 22 | 0 | 0 | - |
| eco00860 Porphyrin and chlorophyll metabolism - Escherichia coli K-12 MG1655 | 24 | 0 | 0 | - |
| eco00910 Nitrogen metabolism - Escherichia coli K-12 MG1655 | 26 | 0 | 0 | - |
| eco00920 Sulfur metabolism - Escherichia coli K-12 MG1655 | 33 | 0 | 1 | 1.02 |
| eco01040 Biosynthesis of unsaturated fatty acids - Escherichia coli K-12 MG1655 | 6 | 0 | 0 | - |
| eco01100 Metabolic pathways - Escherichia coli K-12 MG1655 | 706 | 23 | 13 | 2.85 |
| eco01110 Biosynthesis of secondary metabolites - Escherichia coli K-12 MG1655 | 299 | 15 | 8 | 2.22 |
| eco01120 Microbial metabolism in diverse environments - Escherichia coli K-12 MG1655 | 246 | 14 | 6 | 3.34 |
| eco01130 Biosynthesis of antibiotics - Escherichia coli K-12 MG1655 | 206 | 9 | 4 | 1.97 |
| eco01200 Carbon metabolism - Escherichia coli K-12 MG1655 | 108 | 9 | 3 | 3.18 |
| eco01210 2-Oxocarboxylic acid metabolism - Escherichia coli K-12 MG1655 | 26 | 3 | 0 | 3.18 |
| eco01230 Biosynthesis of amino acids - Escherichia coli K-12 MG1655 | 118 | 6 | 1 | 3.68 |
| eco01501 beta-Lactam resistance - Escherichia coli K-12 MG1655 | 17 | 1 | 0 | 1.03 |
| eco01502 Vancomycin resistance - Escherichia coli K-12 MG1655 | 8 | 0 | 0 | - |
| eco02010 ABC transporters - Escherichia coli K-12 MG1655 | 172 | 2 | 5 | 1.31 |
| eco02020 Two-component system - Escherichia coli K-12 MG1655 | 148 | 0 | 2 | 2.01 |
| eco02024 Quorum sensing - Escherichia coli K-12 MG1655 | 58 | 2 | 0 | 2.04 |
| eco02030 Bacterial chemotaxis - Escherichia coli K-12 MG1655 | 20 | 0 | 1 | 1.03 |
| eco02040 Flagellar assembly - Escherichia coli K-12 MG1655 | 36 | 0 | 1 | 1.01 |
| eco03018 RNA degradation - Escherichia coli K-12 MG1655 | 16 | 0 | 0 | - |
| eco03030 DNA replication - Escherichia coli K-12 MG1655 | 17 | 1 | 0 | 1.03 |
| eco03060 Protein export - Escherichia coli K-12 MG1655 | 18 | 0 | 0 | - |
| eco03070 Bacterial secretion system - Escherichia coli K-12 MG1655 | 30 | 0 | 0 | - |
| eco03410 Base excision repair - Escherichia coli K-12 MG1655 | 14 | 0 | 0 | - |

| | | | | |
|---|---|---|---|---|
| eco03420 Nucleotide excision repair - Escherichia coli K-12 MG1655 | 8 | 0 | 0 | - |
| eco03430 Mismatch repair - Escherichia coli K-12 MG1655 | 22 | 0 | 0 | - |
| eco03440 Homologous recombination - Escherichia coli K-12 MG1655 | 27 | 0 | 0 | - |

**Table S5. Proteins with significant abundance changes in *E. coli* carrying pSU18::*bla*$_{NDM-1}$ versus plasmid only control**

Data provided are raw protein abundance values for three biological replicates (E1-E3). Green, upregulated; red, downregulated > 2 fold.

| Accession | Description | pSU E1 | pSU E2 | pSU E3 | NDM E1 | NDM E2 | NDM E3 | T-test | Fold Change |
|---|---|---|---|---|---|---|---|---|---|
| E5KIY2 | Beta-lactamase NDM-1 | | | | 3.801E9 | 1.402E9 | 1.474E9 | <0.001 | >20 |
| P12995 | Adenosylmethionine-8-amino-7-oxononanoate aminotransferase | | | | 1.004E7 | 3.928E6 | 8.398E7 | <0.001 | >20 |
| P0A8I1 | Putative Holliday junction resolvase | | | | 1.213E7 | 3.244E6 | 1.270E7 | <0.001 | >20 |
| P0AEJ2 | Isochorismate synthase EntC | | | | 6.097E6 | 8.336E6 | 2.845E6 | <0.001 | >20 |
| P26365 | N-acetylmuramoyl-L-alanine amidase AmiB | | | | 1.327E7 | 4.362E6 | 9.108E6 | <0.001 | >20 |
| P31545 | Deferrochelatase/peroxidase EfeB | | | | 4.465E6 | 1.429E6 | 1.471E6 | <0.001 | >20 |
| P77689 | FeS cluster assembly protein SufD | | | | 8.870E6 | 2.600E6 | 3.312E6 | <0.001 | >20 |
| Q46925 | Cysteine desulfurase CsdA | | | | 1.365E7 | 6.250E6 | 3.990E6 | <0.001 | >20 |
| Q47319 | DTW domain-containing protein YfiP | | | | 4.578E6 | 6.224E6 | 1.836E6 | <0.001 | >20 |
| P0C0V0 | Periplasmic serine endoprotease DegP | 2.443E7 | 7.270E6 | 4.277E6 | 1.496E8 | 5.018E7 | 6.648E7 | 0.036 | 7.40 |
| P0A9G6 | Isocitrate lyase | 3.230E7 | 1.361E7 | 6.609E6 | 1.595E8 | 9.265E7 | 9.285E7 | 0.007 | 6.57 |
| P46853 | Uncharacterized oxidoreductase YhhX | 1.150E7 | 2.974E6 | 4.249E6 | 7.068E7 | 2.303E7 | 2.635E7 | 0.048 | 6.41 |
| P0A6A3 | Acetate kinase | 3.418E8 | 7.565E7 | 1.212E8 | 2.003E9 | 6.744E8 | 7.736E8 | 0.045 | 6.41 |
| P61887 | Glucose-1-phosphate thymidylyltransferase 2 | 5.526E7 | 3.539E7 | 1.267E7 | 1.024E8 | 2.224E8 | 3.367E8 | 0.023 | 6.40 |
| P69931 | DnaA regulatory inactivator Hda | 1.434E7 | 1.182E7 | 1.500E7 | 1.448E8 | 5.014E7 | 6.798E7 | 0.032 | 6.39 |
| P07012 | Peptide chain release factor 2 | 7.813E7 | 3.256E7 | 1.126E7 | 3.437E8 | 1.427E8 | 1.811E8 | 0.024 | 5.47 |
| P0A9J0 | Ribonuclease G | 1.140E7 | 1.897E6 | 1.500E6 | 3.858E7 | 2.033E7 | 2.120E7 | 0.016 | 5.41 |
| P76273 | Ribosomal RNA small subunit methyltransferase F | 4.293E6 | 2.205E6 | 4.444E5 | 1.240E7 | 1.084E7 | 1.340E7 | <0.001 | 5.28 |
| P07650 | Thymidine phosphorylase | 1.227E7 | 9.149E6 | 4.538E6 | 6.108E7 | 3.030E7 | 3.875E7 | 0.011 | 5.01 |
| P0A825 | Serine hydroxymethyltransferase | 1.261E8 | 4.022E7 | 3.310E7 | 5.075E8 | 2.172E8 | 2.658E8 | 0.025 | 4.97 |
| P00946 | Mannose-6-phosphate isomerase | 2.647E7 | 4.983E6 | 1.199E7 | 1.123E8 | 4.480E7 | 5.344E7 | 0.033 | 4.84 |
| P25437 | S-(hydroxymethyl)glutathione dehydrogenase | 2.629E7 | 8.545E6 | 1.860E7 | 1.368E8 | 5.648E7 | 6.288E7 | 0.031 | 4.79 |
| Q46812 | Protein SsnA | | 1.685E6 | 1.112E6 | | 6.148E6 | 7.159E6 | 0.006 | 4.76 |
| P0A6K6 | Phosphopentomutase | 6.598E7 | 2.632E7 | 2.749E7 | 2.840E8 | 1.249E8 | 1.542E8 | 0.022 | 4.70 |
| P0A953 | 3-oxoacyl-[acyl-carrier-protein] synthase 1 | 2.490E8 | 8.800E7 | 5.566E7 | 9.216E8 | 4.060E8 | 4.683E8 | 0.027 | 4.57 |
| P0CE48 | Elongation factor Tu 2 | | 1.205E9 | 2.367E9 | | 6.832E9 | 9.316E9 | 0.022 | 4.52 |

| | | | | | | | | | |
|---|---|---|---|---|---|---|---|---|---|
| P0AGG8 | Protein TldD | 2.564E7 | 4.603E6 | 3.769E6 | 8.130E7 | 3.382E7 | 3.551E7 | 0.043 | 4.43 |
| P52612 | Flagellum-specific ATP synthase | 4.394E6 | 8.824E5 | 8.247E5 | 1.326E7 | 7.540E6 | 5.870E6 | 0.027 | 4.37 |
| P24232 | Flavohemoprotein | 4.895E6 | | 2.399E6 | 2.197E7 | 1.064E7 | 1.506E7 | 0.034 | 4.36 |
| P0AF18 | N-acetylglucosamine-6-phosphate deacetylase | 1.018E7 | 2.448E6 | 6.148E6 | 4.021E7 | 1.930E7 | 2.203E7 | 0.020 | 4.34 |
| P0ABZ6 | Chaperone SurA | 5.719E7 | 2.004E7 | 8.115E6 | 1.979E8 | 7.416E7 | 9.661E7 | 0.041 | 4.32 |
| P0AFG6 | Dihydrolipoyllysine-residue succinyltransferase component of 2-oxoglutarate dehydrogenase complex | 8.833E7 | 2.938E7 | 1.904E7 | 2.629E8 | 1.385E8 | 1.826E8 | 0.012 | 4.27 |
| P0ABH7 | Citrate synthase | 7.544E7 | 2.891E7 | 4.860E7 | 2.866E8 | 1.720E8 | 1.945E8 | 0.006 | 4.27 |
| P27306 | Soluble pyridine nucleotide transhydrogenase | 2.069E7 | 8.739E6 | 4.872E6 | 7.317E7 | 3.163E7 | 4.110E7 | 0.025 | 4.25 |
| P40874 | N-methyl-L-tryptophan oxidase | 1.018E7 | 2.476E6 | 3.021E6 | 3.637E7 | 1.401E7 | 1.629E7 | 0.043 | 4.25 |
| P35340 | Alkyl hydroperoxide reductase subunit F | 1.304E8 | 3.040E7 | 3.990E7 | 4.142E8 | 2.218E8 | 2.102E8 | 0.021 | 4.23 |
| P0A6H5 | ATP-dependent protease ATPase subunit HslU | 8.502E7 | 4.034E7 | 2.171E7 | 2.933E8 | 1.269E8 | 1.934E8 | 0.020 | 4.17 |
| P04968 | L-threonine dehydratase biosynthetic IlvA | 2.701E6 | 9.012E5 | 7.431E5 | 6.565E6 | 5.778E6 | 5.687E6 | 0.001 | 4.15 |
| Q46803 | Uncharacterized protein YgeW | 2.537E6 | 4.659E6 | 8.523E6 | 1.557E7 | 2.133E7 | 2.787E7 | 0.007 | 4.12 |
| P23847 | Periplasmic dipeptide transport protein | 6.545E6 | 1.861E6 | 2.051E6 | 2.273E7 | 1.097E7 | 9.305E6 | 0.037 | 4.11 |
| P00934 | Threonine synthase | 5.122E7 | 1.995E7 | 1.270E7 | 1.740E8 | 7.566E7 | 9.274E7 | 0.029 | 4.08 |
| P0A6P9 | Enolase | 8.902E8 | 2.242E8 | 2.395E8 | 2.769E9 | 1.270E9 | 1.461E9 | 0.028 | 4.06 |
| P16095 | L-serine dehydratase 1 | 3.008E7 | 9.030E6 | 6.325E6 | 9.325E7 | 4.562E7 | 4.489E7 | 0.030 | 4.04 |
| P0A799 | Phosphoglycerate kinase | 7.303E8 | 1.685E8 | 3.815E8 | 2.554E9 | 1.184E9 | 1.404E9 | 0.024 | 4.02 |
| C4ZU91 | CinA-like protein | 7.658E6 | 3.316E6 | 3.782E6 | 3.142E7 | 1.250E7 | 1.522E7 | 0.036 | 4.01 |
| P0A6H1 | ATP-dependent Clp protease ATP-binding subunit ClpX | 6.905E7 | 1.441E7 | 1.404E7 | 2.062E8 | 8.434E7 | 9.937E7 | 0.042 | 4.00 |
| P06720 | Alpha-galactosidase | 2.978E7 | 1.716E7 | 2.944E7 | 1.029E8 | 8.626E7 | 1.163E8 | <0.001 | 4.00 |
| P0AFE8 | NADH-quinone oxidoreductase subunit M | 7.780E6 | | 3.215E6 | 1.929E7 | | 2.278E7 | 0.016 | 3.83 |
| P02943 | Maltoporin | 2.622E7 | 2.332E7 | 2.812E7 | 9.107E7 | 9.997E7 | 1.050E8 | <0.001 | 3.81 |
| P0A7D4 | Adenylosuccinate synthetase | 2.528E8 | 9.831E7 | 5.061E7 | 8.056E8 | 3.392E8 | 3.780E8 | 0.041 | 3.79 |
| P0AEA8 | Siroheme synthase | 7.944E6 | 2.440E6 | 1.290E6 | 2.371E7 | 9.025E6 | 1.151E7 | 0.047 | 3.79 |
| P0A850 | Trigger factor | 7.503E8 | 3.729E8 | 1.175E8 | 2.057E9 | 1.139E9 | 1.505E9 | 0.018 | 3.79 |
| P75691 | Uncharacterized zinc-type alcohol dehydrogenase-like protein YahK | 1.691E6 | | 7.799E5 | 6.602E6 | 2.972E6 | 4.436E6 | 0.046 | 3.78 |
| P29745 | Peptidase T | 1.234E7 | 5.268E6 | 2.681E6 | 3.766E7 | 1.787E7 | 2.044E7 | 0.027 | 3.75 |
| P12281 | Molybdopterin molybdenumtransferase | 2.192E7 | 7.484E6 | 7.733E6 | 7.106E7 | 2.846E7 | 3.886E7 | 0.035 | 3.73 |
| P0A6J5 | D-amino acid dehydrogenase small subunit | 1.873E7 | 5.180E6 | 7.241E6 | 5.242E7 | 2.737E7 | 3.583E7 | 0.015 | 3.71 |
| P08178 | Phosphoribosylformylglycinamidine cyclo-ligase | 5.809E7 | 6.249E6 | 3.194E6 | 1.161E8 | 5.479E7 | 7.449E7 | 0.040 | 3.63 |
| P08660 | Lysine-sensitive aspartokinase 3 | 8.585E6 | 2.965E6 | 1.862E6 | 2.015E7 | 1.805E7 | 1.019E7 | 0.017 | 3.61 |

| | | | | | | | | | |
|---|---|---|---|---|---|---|---|---|---|
| P31120 | Phosphoglucosamine mutase | 1.479E8 | 4.518E7 | 3.177E7 | 4.236E8 | 1.828E8 | 2.040E8 | 0.042 | 3.60 |
| P40120 | Glucans biosynthesis protein D | 1.057E7 | 3.582E6 | 2.941E6 | 2.842E7 | 1.904E7 | 1.266E7 | 0.025 | 3.52 |
| P0AB77 | 2-amino-3-ketobutyrate coenzyme A ligase | 8.728E7 | 2.014E7 | 4.175E7 | 2.781E8 | 1.213E8 | 1.189E8 | 0.047 | 3.47 |
| P0A6E9 | ATP-dependent dethiobiotin synthetase BioD 2 | 5.318E6 | 2.363E6 | 8.348E5 | 9.206E6 | 1.031E7 | | 0.014 | 3.44 |
| P0ADR8 | LOG family protein YgdH | 2.329E7 | 6.339E6 | 5.479E6 | 6.155E7 | 2.604E7 | 3.189E7 | 0.043 | 3.40 |
| P0A9T0 | D-3-phosphoglycerate dehydrogenase | 2.025E7 | 9.404E6 | 9.857E6 | 7.263E7 | 2.684E7 | 3.428E7 | 0.049 | 3.39 |
| P0AEX9 | Maltose-binding periplasmic protein | 1.491E8 | 5.087E7 | 2.037E8 | 3.432E8 | 6.207E8 | 3.430E8 | 0.021 | 3.24 |
| P14407 | Fumarate hydratase class I, anaerobic | 2.136E7 | 9.081E6 | 1.735E7 | 7.874E7 | 2.937E7 | 4.635E7 | 0.038 | 3.23 |
| P37349 | PTS-dependent dihydroxyacetone kinase, phosphotransferase subunit DhaM | 2.848E7 | 5.743E6 | 5.332E6 | 6.157E7 | 2.897E7 | 3.585E7 | 0.041 | 3.20 |
| P36938 | Phosphoglucomutase | 7.088E7 | 1.463E7 | 1.699E7 | 1.489E8 | 7.908E7 | 8.200E7 | 0.039 | 3.02 |
| P25553 | Lactaldehyde dehydrogenase | 3.305E7 | 3.818E7 | 2.432E7 | 8.953E7 | 9.402E7 | 1.037E8 | <0.001 | 3.01 |
| P0A6T3 | Galactokinase | 1.612E7 | 4.908E6 | 3.149E7 | 5.126E7 | 4.986E7 | 5.622E7 | 0.006 | 3.00 |
| P0A996 | Anaerobic glycerol-3-phosphate dehydrogenase subunit C | 5.424E6 | 2.616E6 | 1.063E7 | 2.149E7 | 1.442E7 | 1.975E7 | 0.009 | 2.98 |
| P06987 | Histidine biosynthesis bifunctional protein HisB | 1.021E7 | 3.277E6 | 6.872E6 | 2.775E7 | 1.386E7 | 1.637E7 | 0.028 | 2.85 |
| P09832 | Glutamate synthase [NADPH] small chain | 7.354E6 | 1.408E7 | 1.409E6 | 1.723E7 | 2.183E7 | 2.571E7 | 0.017 | 2.84 |
| P0AC33 | Fumarate hydratase class I, aerobic | 3.205E7 | 1.127E7 | 2.510E7 | 9.229E7 | 3.652E7 | 6.072E7 | 0.040 | 2.77 |
| P0C8J8 | D-tagatose-1,6-bisphosphate aldolase subunit GatZ | 6.664E7 | 3.621E7 | 3.504E7 | 1.561E8 | 8.910E7 | 1.269E8 | 0.012 | 2.70 |
| P0A6F3 | Glycerol kinase | 3.242E8 | 2.212E8 | 2.061E8 | 9.012E8 | 5.505E8 | 5.680E8 | 0.012 | 2.69 |
| P68187 | Maltose/maltodextrin import ATP-binding protein MalK | 3.421E7 | 1.680E7 | 4.945E7 | 9.346E7 | 6.909E7 | 1.023E8 | 0.008 | 2.64 |
| P00926 | D-serine dehydratase | 5.756E6 | 6.853E6 | 9.400E6 | 2.278E7 | 1.528E7 | 1.906E7 | 0.004 | 2.60 |
| P27129 | Lipopolysaccharide 1,2-glucosyltransferase | 1.583E7 | 1.099E7 | 5.150E6 | 3.958E7 | 1.853E7 | 2.424E7 | 0.037 | 2.58 |
| P0AC38 | Aspartate ammonia-lyase | 1.328E8 | 9.305E7 | 7.414E7 | 3.216E8 | 2.026E8 | 2.375E8 | 0.009 | 2.54 |
| P0AGM5 | Protein sirB1 | 3.278E6 | 1.858E6 | 1.227E6 | 6.407E6 | 4.154E6 | 5.558E6 | 0.011 | 2.53 |
| P33940 | Probable malate:quinone oxidoreductase | 9.491E6 | 2.415E6 | 8.627E6 | 1.952E7 | 1.487E7 | 1.490E7 | 0.012 | 2.40 |
| P0ACK2 | Putative aga operon transcriptional repressor | 8.034E6 | 4.500E6 | 8.070E6 | 1.882E7 | 1.669E7 | 1.205E7 | 0.009 | 2.31 |
| P07464 | Galactoside O-acetyltransferase | 1.643E7 | 1.402E7 | 1.565E7 | 9.256E6 | 4.675E6 | 6.759E6 | 0.002 | 0.45 |
| P76298 | Flagellar biosynthesis protein FlhA | 1.093E8 | 6.951E7 | 5.097E7 | 1.304E7 | 1.424E6 | 5.776E7 | 0.048 | 0.31 |
| P21893 | Single-stranded-DNA-specific exonuclease RecJ | 1.425E8 | 6.029E7 | 6.092E7 | 2.824E7 | 3.539E6 | 8.487E6 | 0.029 | 0.15 |
| P07109 | Histidine transport ATP-binding protein HisP | 6.274E6 | 3.673E6 | 6.161E5 | | | | <0.001 | <0.05 |
| P07364 | Chemotaxis protein methyltransferase | 3.914E6 | 1.816E6 | 9.167E5 | | | | <0.001 | <0.05 |
| P33931 | Cytochrome c biogenesis ATP-binding export protein CcmA | 3.045E7 | 7.224E6 | 6.214E6 | | | | <0.001 | <0.05 |

**Table S6. Chi Squared analysis of KEGG functional grouping of proteins with significant abundance changes in *E. coli* carrying pSU18::*bla*$_{NDM-1}$ versus plasmid only control.**

| KEGG | Total in Genome | Number with abundance change | random | Chi Sq |
|---|---|---|---|---|
| eco00010 Glycolysis / Gluconeogenesis - Escherichia coli K-12 MG1655 | 43 | 4 | 9 | 2.27 |
| eco00020 Citrate cycle | 27 | 5 | 2 | 1.48 |
| eco00030 Pentose phosphate pathway - Escherichia coli K-12 MG1655 | 30 | 2 | 0 | 2.07 |
| eco00051 Fructose and mannose metabolism - Escherichia coli K-12 MG1655 | 41 | 2 | 2 | 0 |
| eco00052 Galactose metabolism - Escherichia coli K-12 MG1655 | 36 | 4 | 1 | 1.93 |
| eco00053 Ascorbate and aldarate metabolism - Escherichia coli K-12 MG1655 | 13 | 1 | 0 | 1.04 |
| eco00061 Fatty acid biosynthesis - Escherichia coli K-12 MG1655 | 13 | 1 | 1 | 0 |
| eco00071 Fatty acid degradation - Escherichia coli K-12 MG1655 | 15 | 1 | 3 | 1.15 |
| eco00130 Ubiquinone and other terpenoid-quinone biosynthesis - Escherichia coli K-12 MG1655 | 20 | 2 | 0 | 2.11 |
| eco00190 Oxidative phosphorylation - Escherichia coli K-12 MG1655 | 41 | 2 | 2 | 0 |
| eco00220 Arginine biosynthesis - Escherichia coli K-12 MG1655 | 18 | 0 | 1 | 1.03 |
| eco00230 Purine metabolism - Escherichia coli K-12 MG1655 | 89 | 4 | 4 | 0 |
| eco00240 Pyrimidine metabolism - Escherichia coli K-12 MG1655 | 65 | 1 | 2 | 0.34 |
| eco00250 Alanine, aspartate and glutamate metabolism - Escherichia coli K-12 MG1655 | 31 | 3 | 3 | 0 |
| eco00260 Glycine, serine and threonine metabolism - Escherichia coli K-12 MG1655 | 36 | 8 | 0 | 9.00 |
| eco00261 Monobactam biosynthesis - Escherichia coli K-12 MG1655 | 10 | 1 | 1 | 0 |
| eco00270 Cysteine and methionine metabolism - Escherichia coli K-12 MG1655 | 32 | 3 | 1 | 1.07 |
| eco00280 Valine, leucine and isoleucine degradation - Escherichia coli K-12 MG1655 | 11 | 1 | 1 | 0 |
| eco00300 Lysine biosynthesis - Escherichia coli K-12 MG1655 | 15 | 1 | 1 | 0 |
| eco00310 Lysine degradation - Escherichia coli K-12 MG1655 | 12 | 1 | 1 | 0 |
| eco00330 Arginine and proline metabolism - Escherichia coli K-12 MG1655 | 25 | 0 | 2 | 2.08 |
| eco00340 Histidine metabolism - Escherichia coli K-12 MG1655 | 8 | 1 | 0 | 1.07 |

| | | | | |
|---|---|---|---|---|
| eco00350 Tyrosine metabolism - Escherichia coli K-12 MG1655 | 9 | 1 | 1 | 0 |
| eco00360 Phenylalanine metabolism - Escherichia coli K-12 MG1655 | 30 | 1 | 1 | 0 |
| eco00380 Tryptophan metabolism - Escherichia coli K-12 MG1655 | 9 | 0 | 1 | 1.06 |
| eco00400 Phenylalanine, tyrosine and tryptophan biosynthesis - Escherichia coli K-12 MG1655 | 21 | 0 | 2 | 2.10 |
| eco00430 Taurine and hypotaurine metabolism - Escherichia coli K-12 MG1655 | 6 | 1 | 0 | 1.09 |
| eco00450 Selenocompound metabolism - Escherichia coli K-12 MG1655 | 17 | 1 | 0 | 1.03 |
| eco00460 Cyanoamino acid metabolism - Escherichia coli K-12 MG1655 | 6 | 1 | 0 | 1.09 |
| eco00473 D-Alanine metabolism - Escherichia coli K-12 MG1655 | 4 | 0 | 1 | 1.14 |
| eco00500 Starch and sucrose metabolism - Escherichia coli K-12 MG1655 | 36 | 1 | 2 | 0.35 |
| eco00520 Amino sugar and nucleotide sugar metabolism - Escherichia coli K-12 MG1655 | 46 | 5 | 1 | 2.85 |
| eco00521 Streptomycin biosynthesis - Escherichia coli K-12 MG1655 | 9 | 2 | 0 | 2.25 |
| eco00523 Polyketide sugar unit biosynthesis - Escherichia coli K-12 MG1655 | 6 | 1 | 0 | 1.09 |
| eco00525 Acarbose and validamycin biosynthesis - Escherichia coli K-12 MG1655 | 4 | 1 | 0 | 1.14 |
| eco00540 Lipopolysaccharide biosynthesis - Escherichia coli K-12 MG1655 | 31 | 1 | 0 | 1.02 |
| eco00561 Glycerolipid metabolism - Escherichia coli K-12 MG1655 | 12 | 2 | 0 | 2.18 |
| eco00564 Glycerophospholipid metabolism - Escherichia coli K-12 MG1655 | 30 | 1 | 1 | 0 |
| eco00600 Sphingolipid metabolism - Escherichia coli K-12 MG1655 | 3 | 1 | 1 | 0 |
| eco00620 Pyruvate metabolism - Escherichia coli K-12 MG1655 | 51 | 5 | 7 | 0.38 |
| eco00625 Chloroalkane and chloroalkene degradation - Escherichia coli K-12 MG1655 | 4 | 1 | 1 | 0 |
| eco00626 Naphthalene degradation - Escherichia coli K-12 MG1655 | 4 | 1 | 1 | 0 |
| eco00627 Aminobenzoate degradation - Escherichia coli K-12 MG1655 | 8 | 0 | 1 | 1.07 |
| eco00630 Glyoxylate and dicarboxylate metabolism - Escherichia coli K-12 MG1655 | 41 | 4 | 3 | 0.16 |
| eco00640 Propanoate metabolism - Escherichia coli K-12 MG1655 | 38 | 1 | 3 | 1.06 |
| eco00650 Butanoate metabolism - Escherichia coli K-12 MG1655 | 37 | 0 | 1 | 1.01 |
| eco00670 One carbon pool by folate - Escherichia coli K-12 MG1655 | 14 | 1 | 0 | 1.04 |
| eco00680 Methane metabolism - Escherichia coli K-12 MG1655 | 27 | 5 | 2 | 1.48 |

| | | | | |
|---|---|---|---|---|
| eco00750 Vitamin B6 metabolism - Escherichia coli K-12 MG1655 | 9 | 1 | 0 | 1.06 |
| eco00760 Nicotinate and nicotinamide metabolism - Escherichia coli K-12 MG1655 | 22 | 1 | 0 | 1.02 |
| eco00770 Pantothenate and CoA biosynthesis - Escherichia coli K-12 MG1655 | 23 | 0 | 3 | 3.21 |
| eco00780 Biotin metabolism - Escherichia coli K-12 MG1655 | 14 | 3 | 2 | 0.24 |
| eco00790 Folate biosynthesis - Escherichia coli K-12 MG1655 | 22 | 1 | 0 | 1.02 |
| eco00860 Porphyrin and chlorophyll metabolism - Escherichia coli K-12 MG1655 | 24 | 1 | 1 | 0 |
| eco00900 Terpenoid backbone biosynthesis - Escherichia coli K-12 MG1655 | 13 | 0 | 1 | 1.04 |
| eco00910 Nitrogen metabolism - Escherichia coli K-12 MG1655 | 26 | 1 | 1 | 0 |
| eco01040 Biosynthesis of unsaturated fatty acids - Escherichia coli K-12 MG1655 | 6 | 1 | 0 | 1.09 |
| eco01100 Metabolic pathways - Escherichia coli K-12 MG1655 | 706 | 41 | 29 | 2.16 |
| eco01110 Biosynthesis of secondary metabolites - Escherichia coli K-12 MG1655 | 299 | 24 | 15 | 2.22 |
| eco01120 Microbial metabolism in diverse environments - Escherichia coli K-12 MG1655 | 246 | 18 | 15 | 0.29 |
| eco01130 Biosynthesis of antibiotics - Escherichia coli K-12 MG1655 | 206 | 20 | 14 | 1.15 |
| eco01200 Carbon metabolism - Escherichia coli K-12 MG1655 | 108 | 14 | 7 | 2.58 |
| eco01210 2-Oxocarboxylic acid metabolism - Escherichia coli K-12 MG1655 | 26 | 2 | 3 | 0.22 |
| eco01212 Fatty acid metabolism - Escherichia coli K-12 MG1655 | 21 | 1 | 2 | 0.36 |
| eco01220 Degradation of aromatic compounds - Escherichia coli K-12 MG1655 | 16 | 1 | 1 | 0 |
| eco01230 Biosynthesis of amino acids - Escherichia coli K-12 MG1655 | 118 | 12 | 8 | 0.87 |
| eco01501 beta-Lactam resistance - Escherichia coli K-12 MG1655 | 17 | 0 | 3 | 3.29 |
| eco01502 Vancomycin resistance - Escherichia coli K-12 MG1655 | 8 | 0 | 1 | 1.07 |
| eco01503 Cationic antimicrobial peptide (CAMP) resistance - Escherichia coli K-12 MG1655 | 35 | 2 | 5 | 1.43 |
| eco02010 ABC transporters - Escherichia coli K-12 MG1655 | 172 | 6 | 8 | 0.30 |
| eco02020 Two-component system - Escherichia coli K-12 MG1655 | 148 | 3 | 7 | 1.66 |
| eco02024 Quorum sensing - Escherichia coli K-12 MG1655 | 58 | 1 | 0 | 1.01 |
| eco02030 Bacterial chemotaxis - Escherichia coli K-12 MG1655 | 20 | 3 | 1 | 1.11 |
| eco02040 Flagellar assembly - Escherichia coli K-12 MG1655 | 36 | 3 | 0 | 3.13 |

| | | | | |
|---|---|---|---|---|
| eco02060 Phosphotransferase system (PTS) - Escherichia coli K-12 MG1655 | 42 | 0 | 3 | 3.11 |
| eco03018 RNA degradation - Escherichia coli K-12 MG1655 | 16 | 1 | 1 | 0 |
| eco03410 Base excision repair - Escherichia coli K-12 MG1655 | 14 | 1 | 0 | 1.04 |
| eco03430 Mismatch repair - Escherichia coli K-12 MG1655 | 22 | 1 | 0 | 1.02 |
| eco03440 Homologous recombination - Escherichia coli K-12 MG1655 | 27 | 1 | 0 | 1.02 |

**Table S7. Antimicrobial susceptibilities of clinical isolates Ent1/2 and KP3.**

| | **Ent1/2** | **KP3** |
|---|---|---|
| **Amoxicillin/Clavulanate** | R | R |
| **Piperacillin/Tazobactam** | R | R |
| **Cefpodoxime** | R | R |
| **Ceftazidime** | R | R |
| **Ceftriaxone** | R | R |
| **Cefuroxime** | R | R |
| **Ertapenem** | R | R |
| **Meropenem** | I (MIC < 4 mg/L) | R (MIC > 32 mg/L) |
| **Ciprofloxacin** | R | R |
| **Gentamicin** | S | S |
| **Co-Trimoxazole** | R | R |

**Resistant (R), Susceptible (S), Intermediate (I).**

**Table S8. 100% Identical blastn hits to *bla*$_{NDM-1}$ region found in Ent1/2 and KP3.**

| Genbank Accession | Species | Country of Origin |
|---|---|---|
| MG878868.1 | *K. pneumoniae* | CHINA |
| MG878867.1 | *E. coli* | CHINA |
| MG878866.1 | *E. coli* | CHINA |
| MK124610.1 | *Citrobacter werkmanii* | UK |
| CP031216.1 | *E. coli* | UK |
| KP770033.1 | *K. pneumoniae* | PAKISTAN |
| KP770031.1 | *Pseudocitrobacter faecalis* | PAKISTAN |
| KP770029.1 | *P. faecalis* | PAKISTAN |
| KP770030.1 | *E. coli* | PAKISTAN |
| KP770027.1 | *C. freundii* | PAKISTAN |
| KP770023.1 | *E. coli* | PAKISTAN |
| KJ440076.1 | *E. coli* | TAIWAN |
| KJ440075.1 | *K. pneumoniae* | TAIWAN |
| AB769140.1 | *E. coli* | JAPAN |
